# *Vibrio* are a potential source of novel colistin-resistance genes in European coastal environments

**DOI:** 10.1101/2024.10.10.617683

**Authors:** Jamal Saad, Viviane Boulo, David Goudenège, Coralie Broquard, Karl B. Andree, Manon Auguste, Bruno Petton, Yannick Labreuche, Pablo Tris, Dolors Furones, Augusti Gil, Luigi Vezzulli, Gianluca Corno, Andrea Di Cesare, Hugo Koechlin, Emilie Labadie-Lafforgue, Gaelle Courtay, Océane Romatif, Juliette Pouzadoux, Jean-Michel Escoubas, Dominique Munaron, Guillaume M. Charrière, Eve Toulza, Marie-Agnès Travers, Caroline Montagnani, Mathias K. Wegner, Delphine Destoumieux-Garzón

## Abstract

Colistin is a widespread last resort antibiotic for treatment of multidrug-resistant bacteria. The recent worldwide emergence of colistin resistance (Col-R) conferred by *mcr*-1 in human pathogens has raised concern, but the putative sources and reservoirs of novel *mcr* genes in the marine environment remain underexplored. We observed a high prevalence of Col-R, particularly in *Vibrio* isolated from European coastal waters by using a unique stock of specific-pathogen-free (SPF) oysters as a bioaccumulator. The high sequence diversity found in the *mcr/ept*A gene family was geographically structured, particularly for three novel *eptA* gene variants, which were restricted to the Mediterranean (France, Spain) and occurred as a *dgk*A-*ept*A operon controlled by the RstA/RstB two component system. By analyzing 29427 *Vibrionaceae* genome assemblies, we showed that this mechanism of intrinsic resistance is prevalent and specific to the Harveyi clade, which includes strains of *Vibrio parahaemolyticus* and *Vibrio alginolyticus* causing infections in humans. The operon conferred colistin-resistance when transferred to sensitive non-*Vibrio* strains. While *mcr*-and *arn*-based Col-R mechanisms were also identified, the widespread presence of *ept*A gene variants in *Vibrio* suggests they play a key role in intrinsic resistance to colistin. Beyond these ancient *ept*A gene copies having evolved with the *Vibrio* lineage, we also identified mobile *ept*A paralogues that have been recently transferred between and within *Vibrio* clades. This highlights *Vibrio* as a potential source of Col-R mechanisms, emphasizing the need for enhanced surveillance to prevent colistin-resistant infections in coastal areas.

## Introduction

The excessive and inadequate use of antibiotics in human and veterinary medicine has led to the spread of antimicrobial resistance genes (ARGs) by creating selective pressures favoring the development of resistant bacteria^1^. If this problem is not addressed, it is estimated that by 2050, antimicrobial resistant bacteria (ARB) could lead to an annual loss of approximately 10 million lives and limit options for effectively treating bacterial infections^2^.

The antibiotic properties of cationic cyclic antimicrobial peptides belonging to the group of polymyxins (*e.g.* Colistin, polymyxin B) is well known, but the use of these peptides for human therapy is limited due to strong side effects. For this reason, limited resistance is reported in clinical settings, promoting the use of polymyxins as a last resort antibiotic for treatments of multidrug-resistant infections^3–5^. Preservation of long-term effectiveness of polymyxins is thus of primary importance for human health globally. Regrettably, the use of colistin has increased as a growth promoter in poultry and swine farms. This has resulted in the rapid spread of colistin resistance in Gram-negative bacteria that are clinically significant on a global scale^6,7^. The global spread of polymyxin-resistant bacteria in clinical and environmental settings has become a major concern in the treatment of multidrug-resistant pathogens in recent years.

Polymyxins bind to the negatively charged Lipid A component of lipopolysaccharides at the outer membrane of Gram-negative bacteria^5,8^, then they disrupt the structure of the outer membrane, leading to the increase of cell permeability and subsequent cell death^9^. Resistance to polymyxins by Gram-negative bacteria can rely on different mechanisms leading to the reduction of Lipid A negative charges thus decreasing electrostatic interactions with polymyxins^10^. In nature, the most frequently observed colistin resistance genes are *ept*A and *pmr*HFIJFKLM (also referred to as *arn*BCADTEF). These genes are chromosomally encoded and catalyze the addition of phosphoethanolamine (PEtN), or the addition of a 4-amino-4-deoxy-L-arabinose (L-Ara4N) to the phosphate groups of lipids A^11^. In various bacterial genera the activation of the *ept*A and *arn*BCADTEF expression is controlled by the PmrA/B and/or PhoP/Q two-component systems (TCS) (for review see^12^). Additionally, a plasmid-mediated mechanism of resistance to colistin was discovered less than 10 years ago and involves *mcr*-1 (for mobile colistin resistance)^6^. It is a rare example of a recent ARG capture and spread. The emergence and rapid spread of *mcr-*1 among various Gram-negative bacteria has alerted health organizations worldwide (Europe, Asia, North America, and Africa)^13^, and the number of newly reported *mcr* genes is ever-growing since 2015^14^. Furthermore, recent research indicates that the majority of mobile colistin resistance gene variants (*mcr*1-9) originate from environmental bacteria, particularly from aquatic sources^15^. Similar to *eptA, mcr-*1 encodes a phosphoethanolamine transferase (PET) but its expression is not dependent on a regulatory system^10^.

The rapid spread of colistin resistance from environmental sources highlights the urgent need to enhance our understanding of the prevalence and distribution of colistin-resistant bacteria and their associated resistance genes in aquatic ecosystems. This calls for research that emphasizes the interconnectedness of humans, animals, and the environment (*i.e,* the One-Health concept^16,17^). However, the marine environment remains largely unexplored regarding antimicrobial resistance (AMR), especially in Europe. Studying AMR in coastal marine environments is particularly important because (i) coastal systems are highly exposed to human contaminants that may select or co-select for ARGs, (ii) coastal systems are highly interconnected through international trade, which favors the worldwide spread of ARGs, (iii) human populations live along coasts and depend on marine environments as a food source^18^, which increases the risk of transmission, and (iv) coastal waters and sediments, wastewater discharge and marine aquaculture can act as sources/reservoirs of ARGs and have contributed to localized increases in abundance of ARGs^19,20^. Still, the study of AMR in marine waters lags behind in comparison to other environments, and there is a lack of understanding of the role these environments play in the global cycle of AMR.

The marine environment may harbor diverse ARGs, flourishing under human-induced pressures. Almost all known variants of the *mcr* gene that have been reported thus far are from aquatic bacteria from diverse environments ^15^. Among them, aquatic bacteria from the *Shewanella* genus are considered as a source of *mcr-*4^15^. Moreover, *mcr-*1 was found recently in colistin-resistant bacteria in marine coastal waters from Norway and Croatia, which may constitute reservoirs^21,22^. Some recent studies have also shown the role of variants of the *ept*A gene in colistin resistance in strains of *Vibrio cholerae*^23^*, Vibrio parahaemolyticus*^24^ *Vibrio vulnificus*^25^ and *Vibrio fisheri* (*Aliivibrio fisheri*)^26^. Given the limited data available on the frequency and distribution of colistin-resistant bacteria in marine environments, along with the evidence of colistin-resistant *Vibrio* species with pathogenic potential in humans and marine fauna^27^, there is an urgent need to increase our understanding of the medical and ecological significance of this phenomenon.

To further evaluate the potential role of the coastal environment in the emergence and circulation of colistin resistance genes, we determined the prevalence of *mcr*/*ept*A genes in culturable marine bacteria in coastal waters across Europe, as well as in *Vibrionaceae* in general. We targeted three different European areas that are heavily impacted by anthropogenic pollution from oyster farming and tourism: Sylt (Germany), the Ebro Delta (Spain), and Thau Lagoon (France). We characterized the prevalence, diversity, and distribution of *mcr/ept*A by using a unique stock of SPF-oysters as bioaccumulators that were incubated in all sites for tracking AMR, as they concentrate bacteria from the environment by filter feeding^28^. Our data revealed an unexpected diversity of *ept*A in *Vibrio*, with a clearly structured geographic distribution of *ept*A variants across Europe. Functional genetic experiments were used to demonstrate the mechanisms of colistin resistance conferred by the newly discovered *eptA* gene variants from the *Vibrio* Harveyi clade. The discovery of highly diverse and prevalent mechanisms of colistin resistance as well as as evidences of recent mobilization in *Vibrionaceae* highlights the underestimated risk of the emergence of colistin resistance in European coastal environments and warrants further investigation.

## Results

### Colistin-resistant bacteria are abundant in oysters in European coastal environments

To estimate the prevalence of colistin resistance (Col-R) in culturable marine bacteria from European coastal ecosystems, we immersed SPF-oysters in three sites either containing natural beds (Sylt, Germany), or oyster farms (Ebro, Spain and Thau, France). After 2-3 weeks, marine bacteria isolated from the SPF-oysters, on either marine agar or Thiosulfate Citrate Bile Salts Sucrose (TCBS) agar at 20°C or 37°C, were tested for Col-R using a microtiter plate assay. Bacteria isolated on marine agar showed frequent resistance to colistin with a minimal inhibitory concentration (MIC) > 5 µg/ml in Zobell medium, which mimics the composition of seawater (Table 1). Out of 87 bacterial isolates from Thau (France), 28 (32.1%) were Col-R. Most of these Col-R isolates (27/28) were isolated at 37°C. In Ebro (Spain), 17/140 isolates were Col-R (12.1%) and most of them (11/17) were isolated at 37°C. In Sylt (Germany), 37/51 isolates were Col-R (72.5%), and most of them (23/37) were isolated at 37°C (Table 1, Fig. S1). Col-R phenotypes were even more prevalent among bacteria isolated on TCBS (selective for *Vibrio*). Out of the 52 isolates obtained on TCBS from Thau samples, 16 (30.8%) were Col-R. Among them, 11/16 were isolated at 20°C and 5/16 were isolated at 37°C. In Ebro, 11 out of 44 isolates (25%) were Col-R. Among them, 5/11 were isolated at 20°C and 6/11 were isolated at 37°C. In Sylt, most isolates (41/45; 91.1%) were Col-R; all of them were isolated at 20°C (no bacterial isolates at 37°C) (Table 1). Overall, our sampling highlighted a remarkable prevalence of colistin resistant isolates in culturable bacteria accumulating in oysters immersed in European coastal waters, particularly in North Germany (general linear model, *p =* 1.5 x 10^-5^). In France and Spain, the percentage was particularly high in bacteria isolated at 37°C (*p=*3.8 x 10^-5^). There was also a clear effect of the isolation on TCBS medium (*p=* 7.7x 10^-3^), suggesting higher Col-R prevalence in *Vibrio* (Fig. S1).

**Table 1.** Colistin-resistant (Col-R*) bacterial isolates** across European oyster farms.

| Sampling site | Isolation<br>medium | No. isolates<br>(20°C; 37°C) | No. Col-R isolates<br>(20°C; 37°C) | % Col-R isolates |
| --- | --- | --- | --- | --- |
| THAU | Marine agar | 87 (41; 46) | 28 (1; 27) | 32.1 |
| EBRO | Marine agar | 140 (96; 44) | 17 (6;11) | 12.1 |
| SYLT | Marine agar | 51 (19;32) | 37 (14;23) | 72.5 |
| THAU | TCBS | 52 (47; 5) | 16 (11; 5) | 30.8 |
| EBRO | TCBS | 44 (28; 16) | 11 (5; 6) | 25.0 |
| SYLT | TBCS | 45 (45; 0) | 41 (41; 0) | 91.1 |

\* Col-R, MIC > 5 µg/ml in Zobell medium
\*\* bacteria isolated on TCBS or Marine agar medium

### High diversity of *mcr*/*ept*A genes in oysters in European coastal systems

To capture the diversity of *mcr*/*ept*A resistance genes circulating in culturable bacteria from European marine coastal systems, pool-sequencing was performed on a total of 394 bacterial isolates pooled by site and isolation conditions (culture medium, temperature) with an average number of 4.10^7^ reads per pool (99 to 149 strains per sampling site; see Table S1). A total of 53 complete unique nucleotide sequences related to *mcr*/*ept*A were found in the contigs from the pool-sequencing after Prokka annotations (Table S1 and S2). Among them, 31 and 22 sequences were carried by bacteria isolated on either marine agar or TCBS, respectively (Table S1). The diversity of the Mcr/EptA amino acid sequences was studied by reconstructing their molecular phylogeny. Amino acid sequences deduced from the pool sequencing were compared to the 104 Mcr-1 to -10 amino acid sequences present in the CARD database. The whole set of sequences was also used to identify Mcr/EptA sequences encoded in the 29427 *Vibrionaceae* genome assemblies found in GenBank. A total of 27921/29427 genomes encoded at least one Mcr/EptA, representing 4075 distinct amino acid sequences among 31813 protein hits, which were included in the analysis. EptA and Mcr sequences could clearly be distinguished based on the molecular phylogeny of their deduced amino acid sequences (Fig. 1).

**Fig. 1.**
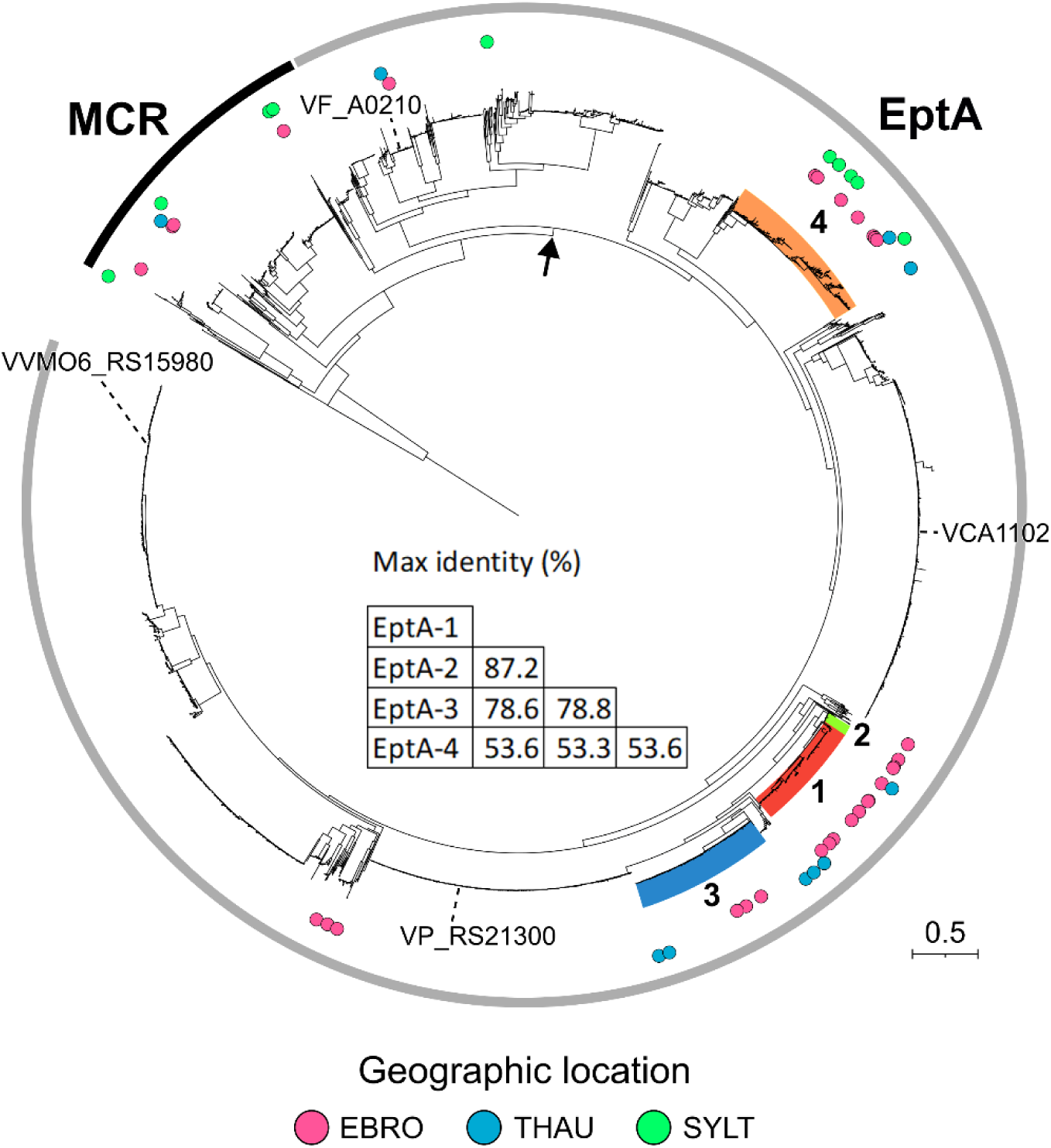
Clustered distribution of *mcr/eptA* gene variants across European coastal environments. The detected EptA and Mcr variants from the present study are included in a phylogenetic tree together with 4075 distinct Mcr/EptA sequences found in 27921 *Vibrionaceae* assemblies carrying *mcr/ept*A genes. Sequences used as reference are Mcr-1 to 10 sequences found in the CARD database as well as 4 EptA sequences from functionally characterized EptA variants (VP_RS21300, VVMO_RS15980, VCA1102 and VF_A0210). A phylogenetic tree generated from deduced EptA/Mcr amino acid sequences was constructed using fasttree with LG model and visualization was done using iTOL. Corresponding protein sequences were obtained from the pool-sequencing of bacteria isolated both on Marine agar and TCBS medium in three European regions. See Fig. S2 for an illustration including conditions of isolation and read counts. The node that separates Mcr (outer black arc) from EptA sequences (outer grey arc) is indicated by an arrow. Numbers refer to the EptA-1(red), EptA-2 (green), EptA-3 (blue) and EptA-4 (orange) variants described in the present study. Maximum identity between newly identified EptA protein variants is displayed. The geographic origin of each sequence is represented in by dots colored pink (Ebro), green (Sylt), and blue (Thau).

The EptA clade gathered a highly diversified set of sequences from *Vibrionaceae*, including the four previously characterized EptA from *V. cholerae*^23^*, V. parahaemolyticus*^24^*, V. vulnificus*^25^ and *V. fisheri*^26^ (Fig. 1). Most of the sequences from the pool-sequencing (42/53) were assigned to EptA (83.87% and 77.27% of the sequences from bacteria isolated on marine agar and on *Vibrio*-selective TCBS medium, respectively) (Fig. S2, Tables S3). They were only found in *Vibrionaceae* (in genera *Vibrio*, *Photobacterium, Allivibrio)* based on protein sequence similarity (more than 90% amino acid sequence identity over their full-length sequence) (Table S4). Only a limited number of sequences (9/53) were assigned to Mcr (Fig. 1, Table S4); they were found in more diverse genera including *Vibrio, Photobacterium, Shewanella,* and *Pseudoalteromonas* (Fig. S2). These sequences encoding Mcr protein variants were previously unreported and the majority clustered with Mcr-4 (Fig. S2). They were far less abundant than sequences encoding EptA protein variants, both in terms of sequence diversity and read counts (Fig 1, Fig. S2).

### EptA variants specific of the Harveyi and Splendidus clades are dominant in oysters

Four predicted EptA proteins (EptA-1 to -4) encoded by unique nucleotide sequences were dominant in oysters, both in terms of sequence diversity (Fig. 1) and read counts (Fig. S2). EptA-1 to -4 sequences harbored the catalytic threonine conserved in EptA orthologs functionally characterized in *Vibrionaceae* (Fig. S3) as well as in EptA/Mcr proteins from *Enterobacteriaceae*^23^. EptA-1 to -3 clustered separately from EptA-4 in the Mcr/EptA phylogeny (Fig. 1). Compared to previously characterized EptA variants, EptA-1 to -3 amino acid sequences showed the maximum identity 80.9-83.6 % with VP_RS21300 from *V. parahaemolyticus* and 69.9-72.9 % with VVMO6_RS15980 from *V. vulnificus*. Only 58.2-58.7 % maximum identity was found with VCA1102 from *V. cholerae* and 42.5-44.5 % with VF_A0210 from *V. fisheri*. EptA-4 was far more divergent with only 43.6 %, 55.2 %, 54.6 % and 56.6 % maximum identity with VF_A0210, VCA1102, VP_RS21300 and VVMO6_RS15980 respectively (Table S4).

EptA-1 (encoded by 10 unique nucleotide sequences), EptA-2 (3 unique nucleotide sequences) and EptA-3 (encoded by 5 unique nucleotide sequences) clustered together on the Mcr/EptA phylogenetic tree (Fig. 1). EptA-1 and EptA-2 showed the highest maximum identity (87.2%), compared to a maximum of 78.6-78.8% between EptA-1/2 and EptA-3 (Fig. 1, Table S4). Moreover, their encoding genes shared a common, but previously undescribed genomic environment, consisting of five conserved genes: *rst*A-*rst*B-Glycine zipper family protein-*dgk*A-*ept*A (Fig. 2B, Table S3). This genomic environment was specific to the Harveyi clade when considering the four genes upstream and downstream of the *mcr/eptA* sequences in a *Vibrionaceae* phylogenetic tree, which was constructed from 23,642 genome assemblies with sufficient quality to be included in the MLSA (Fig. 2A, Dataset 1). While the *rst*A-*rst*B-Glycine zipper family protein-*dgk*A-*ept*A genomic environment was a clear indicator of the assignation to the Harveyi clade, EptA polymorphism appeared to have followed differentiation between species: EptA-1 was found in the species *V. harveyi, V. campbelli, V. jasicida, V. owensii,* and *V. hyugaensis;* EptA-2 was found in *V. rotiferianus*; and EptA-3 was found in *V. alginolyticus* (Fig. 2A). In addition, a very large cluster of EptA sequences similar to VP_RS21300 (reference sequence not sampled in our pool sequencing), which clustered with EptA-1 to -3 on the Mcr/EptA phylogenetic tree (Fig. 1), was found in *V. parahaemolyticus* (Fig. 2A). Finally, an additional EptA sequence found in our pool sequencing, which clustered close to EptA-3 (Fig. 1) and shared the same genomic environment. It matched with *V. alfacsiensis,* also belonging to the Harveyi clade (Table S3).

**Fig. 2.**
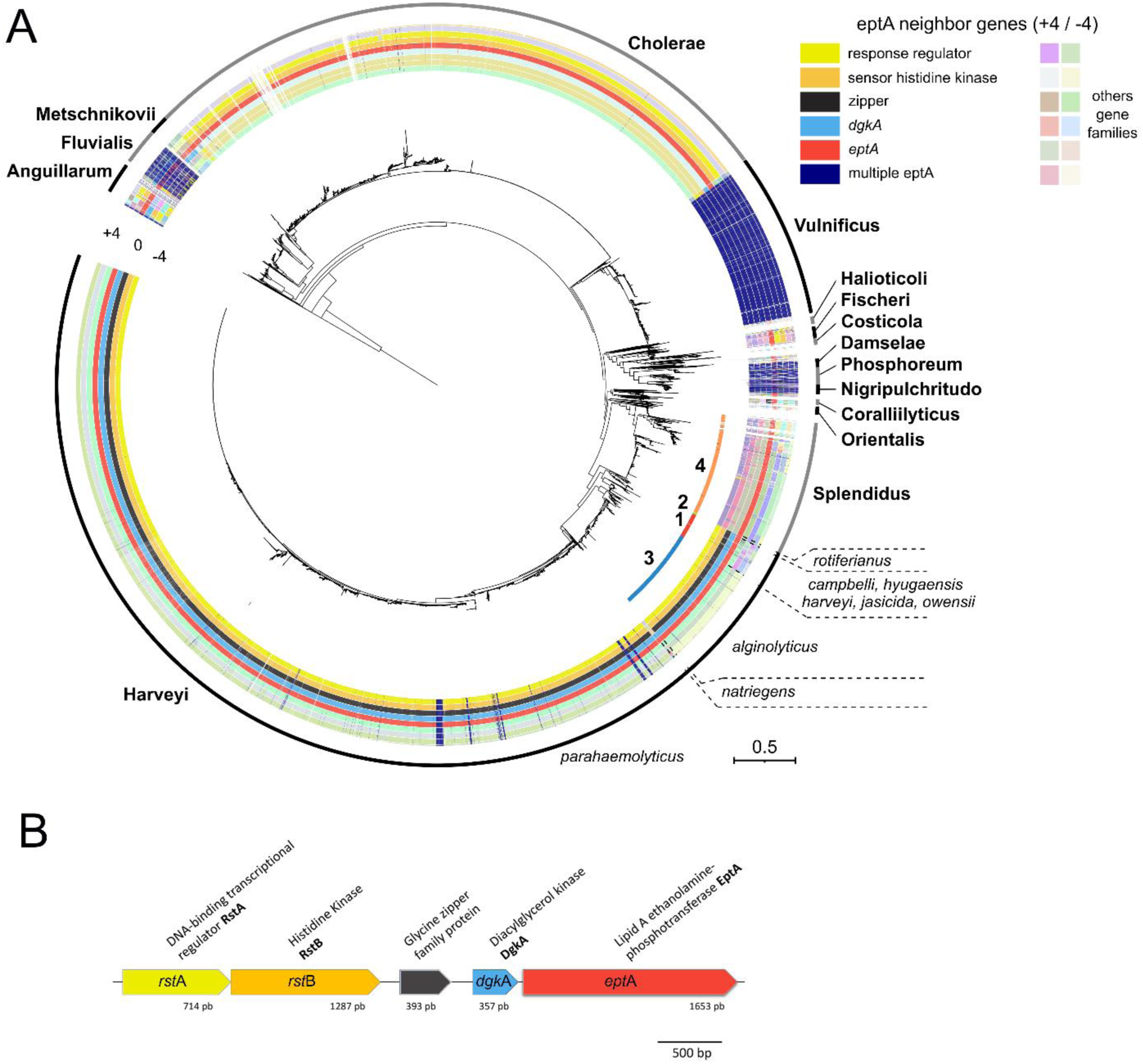
The genomic environment of *ept*A gene variants has evolved with the phylogeny of *Vibrionaceae*. **(A)** Distribution of *mcr/ept*A genomic environments (-4/+4 genes) along an MLST of *Vibrionaceae*. The *Vibrionaceae* phylogenetic tree was based on 8 polymorphic genes *fts*Z, *gap*A, *mre*B, *rpo*A, *top*A, *gyr*B, *pyr*H, *rec*A according to *Vibrio* Clade 3.0^60^. Numbers on the inner circle indicate the phylogenetic positioning of EptA-1, EptA-2, EptA-3, and EptA-4 variants identified in the present study (Table S2). Outer circles show genes in the genomic environment of *mcr/eptA* genes (red) with the following color code: response regulator (yellow), sensor histidine kinase (orange), *dgk*A (light blue), glycine zipper family protein (grey). Other conserved gene families, identified through diamond clustering, are shown using different light colors. When > 1 *mcr/ept*A gene copy were found in *Vibrio* genomes, their genomic environments were not displayed (dark blue). **(B)** Newly discovered genetic environment in the Harveyi clade. A *rst*A*/rst*B (response regulator/histidine kinase) two component system is located upstream *dkgA* and *eptA*. A gene encoding a glycine zipper family protein separates *rst*A*-rst*B from *dgk*A*-ept*A. The figure is based on an *ept*A-1 sequence.

EptA-4 (encoded by 14 unique nucleotide sequences) was the other abundant EptA variant found in our pool sequencing (Fig. 1), but it was a more divergent sequence with only 53.3-53.6% maximum identity with EptA-1 to -3 (Fig. 1). Moreover, the *ept*A-4 genomic environment differed completely, lacking both *dgk*A and the *rstA/rstB* two-component regulatory system (Fig. 2, Table S3). Within the *Vibrionaceae* phylogeny EptA-4 was specific to the Splendidus clade (Fig. 2A).

Two EptA sequences almost identical to *V. fisheri* VF_A0210 (> 98.7 % identity) were carried by bacterial isolates from our European sampling and clustered apart from the newly discovered EptA-1 to -4 protein variants (Fig. 1) (Table S4, Fig.1). The remaining sequences did not share any significant conservation of the sequence/synteny with other EptA variants and were present in sequences of *Vibrio, Allivibrio* and *Photobacterium* outside the Harveyi and Splendidus clades, as well as in the species *Halomonas* (Table S3).

### Existence of both ancient and mobile *ept*A paralogues in the Harveyi clade

In order to determine the potential risk of *ept*A gene transfer from *Vibrionaceae*, we analyzed the genomic environment of *ept*A genes in 24,243 genomes. The majority of *ept*A genes were single copy (Fig. S4) (19,118 / 24,243 genomes), located on chromosomes and we did not detect any insertion sequences, phage fragments, or plasmid fragments in the proximity of - 4/+4 genes around these single copy *ept*A genes, supporting the ancient acquisition of *ept*A genes in *Vibrio*. However, we also identified 2,543 genomes carrying 2 copies and 13 genomes carrying 3 copies of *ept*A (Fig. S4). Some of these additional copies showed sign of recent mobilization as indicated by the presence of transposases/integrases near 137 *ept*A genes from *Vibrionaceae* (0.57 % of *ept*A sequences) and 22 *ept*A genes predicted to be carried on a plasmid (Dataset1). In species of the Harveyi clade, the conserved *ept*A copy (*e.g. ept*A-1, -2 and -3) occurred in a *rst*A-*rst*B-glycine zipper-*dgk*A*-ept*A genomic environment with no trace of mobile genetic elements in its close vincinity (Fig. 2-3). Some Harveyi species such as *V. parahaemolyticus* also harbored a second copy of *ept*A in a distinct genomic environment (*cyt*B-*pep*SY-*ept*A-*dgk*A) with no evidence of mobility (Fig. 3). Remarkably, additional *ept*A paralogues with transposases or integrases at their close vincinity (-4/+4 genes) were found in a *cyt*B*-ept*A*-dgk*A genomic environment (Fig. 3). They showed much closer similarity with *ept*A genes from other *Vibrio* species (*V. cholerae, V. anguillarum*) than with the conserved *ept*A copy from the Harveyi clade (Fig. 3). These results strongly suggest a recent mobilization of *ept*A paralogues between the Harveyi, Cholerae and Anguillarum clades. Similar events of putative *ept*A mobility were also evidenced within the clade Fluvialis where putative mobile paralogues were found in a PAP2-*dgk*A-*ept*A genomic environment (Fig. 3). Altogether, our results highlight the existence in *Vibrionaceae* of an ancient *ept*A gene acquisition having evolved with the *Vibrionaceae* lineage, and largely conserved across *Vibrionaceae* species, as well as rarer and more recent *ept*A gene mobilizations within and between *Vibrio* clades.

**Fig. 3.**
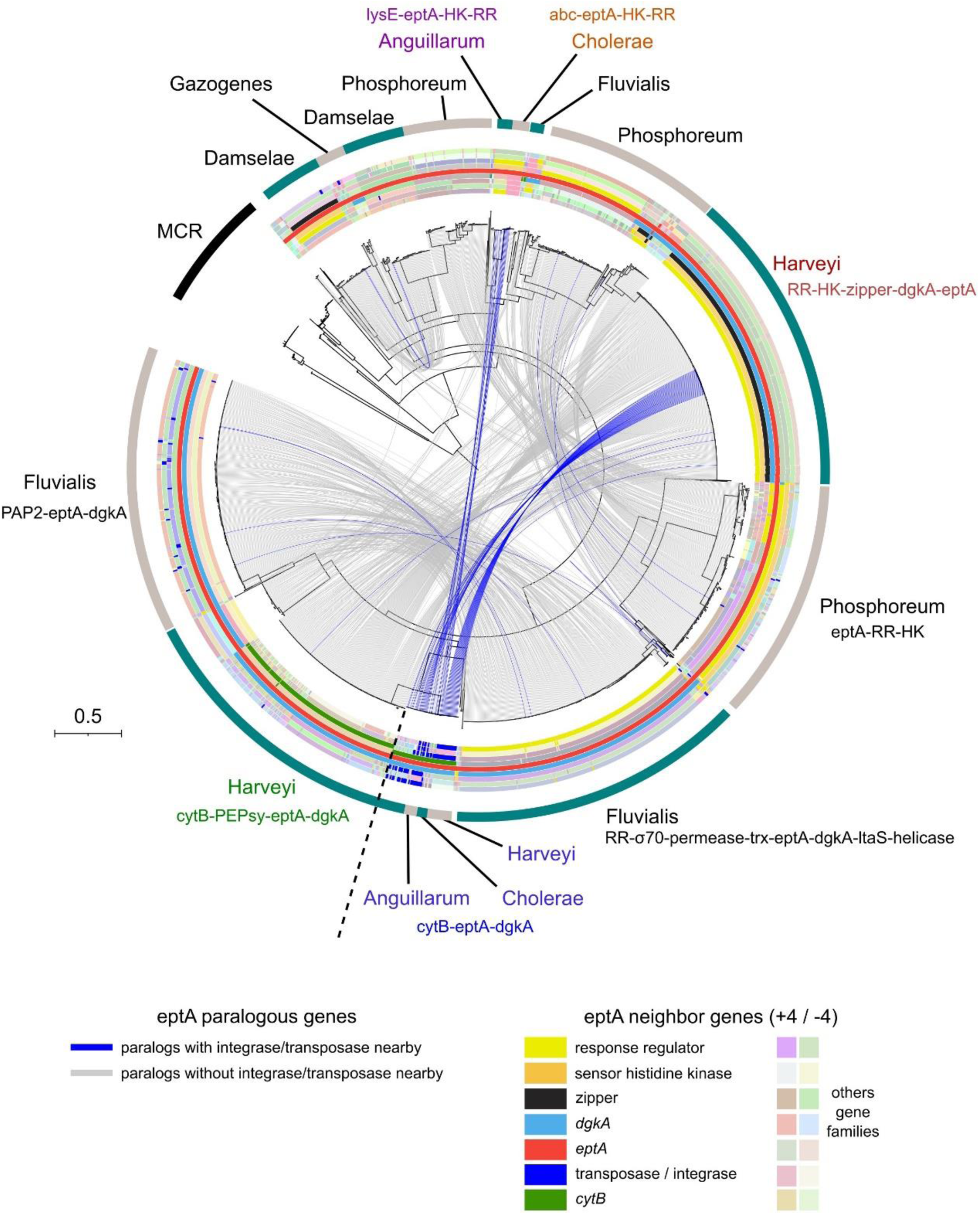
Evidence of gene mobilization in *ept*A paralogues from *Vibrionaceae*. Phylogeny of *ept*A paralogues for genomes containing multiple *ept*A genes, focusing on the distribution of *ept*A genomic environments (-4/+4 genes). The *ept*A phylogenetic tree was constructed using FastTree with the LG model, and visualization was performed with iTOL. To simplify the figure*, V. vulnificus*, which contains two conserved *ept*A paralogues (see Fig. S4) was not included. All paralogues are connected by inner grey lines, while inner blue lines indicate paralogues associated with integrase/transposase in the -4/+4 environ-ments. The outer circles represent genes in the genomic environment of *ept*A genes (in red), with the following color coding: response regulators (yellow), sensor histidine ki-nases (orange), *dgk*A (light blue), glycine zipper family proteins (grey), transposases/inte-grases (dark blue), and cytochrome B family proteins (dark green). Other conserved gene families identified through diamond clustering are depicted in various light colors. The outer circle indicates the Sawabe clade^60^ and provides an environmental summary.

### EptA variants show distinct geographic distribution in Europe

The most common EptA protein variants identified in our sampling were ancient copies conserved in the Harveyi and Splendidus clades and clearly clustered according to geographic locations (Fig. 1). Specifically, a major Mediterranean cluster dominated by EptA-1-2-3 (Harveyi clade) was observed, which includes sequences obtained from EBRO and THAU. It drives the significant geographic structuring between the Mediterranean and North Sea sites (PERMANOVA all sites: F_2,68_ = 3.032, P < 0.001, THAU vs. SYLT: F_1,34_ = 3.108, P = 0.007, EBRO vs. SYLT: F_1,50_ = 4.799, P < 0.001, THAU vs. EBRO: F_1,52_ = 0.900, P = 0.513). Within this Mediterranean cluster EptA-1 and EptA-3 were the more frequently detected variants (Fig. S2). Outside this Mediterranean cluster, EptA-4 from the Splendidus clade was the other abundant EptA variant (Fig.1, Fig. S2). However, geographical structuring within EptA-4 was much weaker and no significant association with a given environment was found (PERMANOVA F_2,16_ = 2.888, P = 0.076) (Fig. 1).

### EptA variants from Harveyi but not Splendidus clade are associated with intrinsic colistin resistance

To investigate the potential role of EptA variants from the Harveyi clade in colistin resistance, we conducted whole genome sequencing (WGS) on nine randomly chosen strains isolated from Thau displaying resistance to colistin. Among the selected strains, three strains from the Harveyi clade, *Vibrio jasicida* TH21_20A_OE8, *Vibrio owensii* TH21_37_OE7, and *Vibrio* sp. TH21_20A_OB7, possessed the *ept*A-1 gene. Three other strains also affiliated to the Harveyi clade, *V. alginolyticus* TH21_37A_OE12, *V. alginolyticus* TH21_37_OE9, and *V. alginolyticus* TH21_37A_OE10, carried the *ept*A-3 gene (Fig. 3, Table S5). To enrich our collection, we explored genomes of *Vibrio* strains collected over the past ten years in French oyster farms. We identified *ept*A-1 and *ept*A-2 in four and two strains of the Harveyi clade, respectively, collected in Thau. We also included two *Vibrio* strains with sequenced genomes and known pathogenic potential. The first was *V. parahaemolyticus* strain IFVp22^29^: it harbors the *ept*A gene variant from the species in the Harveyi *rst*A-*rst*B-glycine zipper-*dgk*A-*ept*A genomic environment. The second, from outside the Harveyi clade, was the zoonotic *V. vulnificus* CECT4999^30^, which harbors a distinct *ept*A gene variant in a *car*R*-caS*-*dgk*A-*ept*A genomic environment, where *car*RS (also known as *vpr*AB) is homologous to *rst*AB. Regarding strains carrying the *ept*A*-*4 gene variant from the Splendidus clade, we found the gene variant in four strains collected in Brest (French Brittany), affiliated with *V. splendidus* and *V. crassostreae* (Fig. 4, Table S5).

**Fig. 4.**
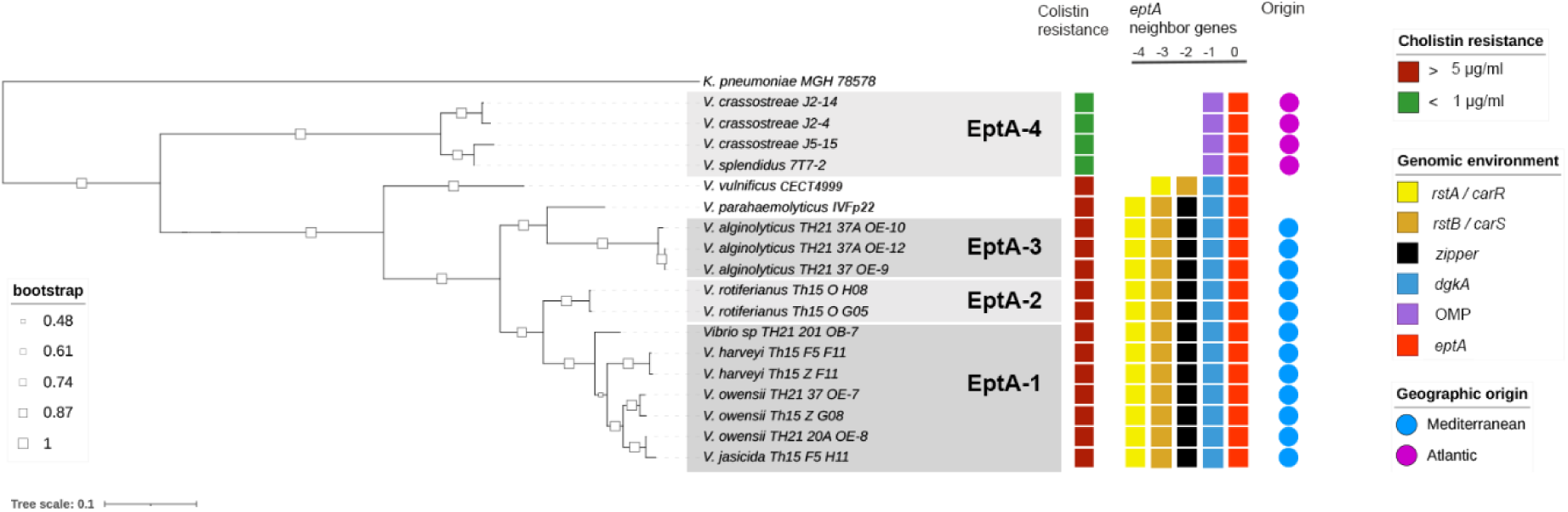
Colistin resistance correlates with EptA polymorphism and *ept*A genomic environment. A phylogenetic tree of EptA amino acid sequences was generated through maximum likelihood analysis of deduced EptA/Mcr amino acid sequences using PhyML v-3.0 (https://ngphylogeny.fr/tools/) with the WAG model and 100 bootstrap runs. Red and green empty squares indicate strains resistant to 5 µg/ml or susceptible to 1 µg/ml colistin, respectively, as phenotyped in Zobell medium. Colored plain squares indicate conserved genes present at the vicinity of *ept*A (red): *rst*A /*car*R (yellow), *rst*B / *car*S (orange), *dgk*A (light blue), glycine zipper family protein (grey), *opa* (purple). Numbers indicate the position of the upstream genes relative to EptA.

Phenotypes of Col-R correlated perfectly with *ept*A variants and their associated genomic environment, as observed by comparing strain phenotypes and single genome sequences for the 16 strains harboring *ept*A-1 to -4 genes (isolated here from oysters) and the two additional pathogenic *Vibrio* strains (Fig. 4). Thus, *Vibrio* strains with *ept*A-1 to -3 were resistant to > 5 µg/ml colistin in Zobell medium, similar to the pathogenic strains *V. parahaemolyticus* IFVp22 and *V. vulnificus* CECT4999. In contrast, strains carrying *ept*A*-*4 were susceptible to colistin at a concentration £ 1 µg/ml in the same conditions (Table S5, Fig. 4), suggesting that *ept*A-1 to *-*3, but not *ept*A-4 confer resistance to colistin. Moreover, the *dgk*A gene is consistently found adjacent to *ept*A in resistant strains (Fig. 4) and in the Harveyi and Vulnificus clades in general (Fig. 4). In contrast, the *dgk*A gene is absent from *ept*A genomic environment in the 4 colistin-susceptible strains harboring *ept*A-4 (Fig. 4) and in the Splendidus clade in general (Fig 2A). In strains of the Harveyi clade, we also noted the conservation of the *rst*A-*rst*B two component signal transduction system (*car*R-*car*S in the Vulnificus clade), located 1084-bp upstream of the *ept*A gene and 706-bp of the *dgk*A gene (Fig. 2 & 4). A CDS encoding a potential glycine zipper protein separated *rst*A-*rst*B from the *dgk*A*-ept*A operon in the Harveyi clade (Fig 2).

### *dgk*A is required for *ept*A-mediated colistin resistance in the Harveyi clade

The co-occurrence of *dgk*A and *ept*A in resistant strains of the Harveyi clade prompted us to test their role in resistance to colistin. Gain of function assays were performed by cloning genes of interest into the pBAD-TOPO expression vector used to transform a colistin-susceptible strain of *E. coli.* Cloning was conducted in *E. coli* TOPO10. Basically, we cloned the naturally occurring *dgk*A-*ept*A-1 and *ept*A-4 under the control of a pBAD promotor. In addition, we cloned *ept*A-1 alone and *dgk*A (from *eptA*-1) alone under the control of pBAD. Among these four constructs, only *dgkA-eptA-1* increased *E. coli* TOPO10 resistance to colistin (MIC > 16 µg/ml in Zobell medium) upon promotor induction. In contrast, the other three constructs did not impact the resistance of *E. coli* TOPO10 to colistin (MIC = 0.25 µg/ml in Zobell medium) (Table 2). The result was confirmed in Muller-Hinton (MHCA) medium (European Committee on Antimicrobial Susceptibility Testing, EUCAST conditions) where only *ept*A*-dgk*A-1 expression increased the MIC of colistin from 0.5 to 4 µg/ml (Table 2). This demonstrates that neither *ept*A*-*1 nor *ept*A*-*4 alone can confer resistance to colistin. Instead, it shows that *dgk*A and *ept*A-1 act together to confer Col-R, in agreement with the conserved genomic environment of *ept*A gene variants 1, 2 and 3 isolated from Mediterranean coastal environments.

**Table 2.** Role of *ept*A variants, *dgk*A and *rst*A in resistance to colistin. MICs were determined by the microdilution assay in the range of 0.125 - 16 μM colistin.

| Strain | Genotype | MIC (μg/ml) | |
| --- | --- | --- | --- |
|  |  | MHCA | Zobell |
| *E. coli* TOP10 | wild type | 0.5 | 0.25 |
| *E. coli* IHPE 10466 | TOP10 pBAD-TOPO-*eptA*-1 * | 0.5 | 0.25 |
| *E. coli* IHPE 10471 | TOP10 pBAD-TOPO-*dgkA* * | 0.5 | 0.25 |
| *E. coli* IHPE 10469 | TOP10 pBAD-TOPO-*dgkA-eptA*-1 * | **4** | **> 16** |
| *E. coli* IHPE 10467 | TOP10 pBAD-TOPO-*eptA*-4 ** | 0.5 | 0.25 |
| *V. harveyi* IHPE 390 | wild type | **8** | **> 16** |
| (Th15_F5-F11) | *rstA-rstB*-glycine zipper-*dgkA-eptA*-1 |  |  |
| *V. harveyi* IHPE 10451 | Th15_F5-F11 Δ*rstA* | 0.5 | 2 |
| *V. harveyi* IHPE 10453 | Th15_F5-F11 Δ*rstA* pMRB-*rstA* | **8** | **> 16** |
| *V. harveyi* IHPE 10452 | Th15_F5-F11 Δ*rstA* pMRB-*gfp* | 0.5 | 2 |
| *E. coli* O6 |  | 1 | 1 |
| (ATCC 25922) |  |  |  |

\* cloned from *V. owensii* Th15\_Z\_G08 (IHPE 303)
\*\* cloned from *V. tasmaniensis* 7T7\_2 (IHPE 20)

### *rst*A/B controls colistin resistance mediated by *dgk*A*-ept*A in the Harveyi clade

We finally tested the potential role of the conserved *rst*A/B two-component system in the expression of colistin resistance in Mediterranean *Vibrio* strains resistant to colistin. For that, we performed *rst*A deletion by allelic exchange in *V. harveyi* strain Th15_F5-F11, which harbors the conserved *rst*A-*rst*B-glycine zipper-*dgk*A*-ept*A1 gene cluster and resists to 16 µg/ml colistin in Zobell medium (Table 2). The Col-R phenotype was significantly compromised in the *rst*A deletion mutant, since growth was fully inhibited in the presence of 2 µg/ml colistin (Table 2). Moreover, complementation with the pMRB-*rst*A plasmid was sufficient to restore full growth at 16 µg/ml Colistin (Table 2). Similarly, in EUCAST conditions, the resistance phenotype of the wild-type Th15_F5-F11 (MIC = 8 µg/ml in MHCA medium), changed to susceptible in the *rst*A deletion mutant (MIC = 0.25 µg/ml in MHCA medium) and was fully restored in the *rst*A complemented strain, but not in a complementation control with *gfp* (Table 2). To demonstrate that the loss of the resistance phenotype is due to an altered expression of *dgk*A and *ept*A in the *rst*A deletion mutant, we quantified the transcripts of *dgk*A and *ept*A by RT-qPCR in the wild-type, mutant, and complemented background. As anticipated, the expression of *dgk*A and *ept*A genes exhibited a significant decrease in the *rst*A deletion mutant (Fig. 5). On the other hand, expression levels were not significantly different between the wild-type *V. harveyi* Th15_F5-F11 strain and its isogenic *rst*A mutant complemented with pMRB-*rst*A (Fig. 5). Such a functional complementation was not observed with the pMRB-*gfp* control plasmid. This demonstrates that *rst*A/B controls the Col-R phenotype of *V. harveyi* Th15_F5-F11 through the expression of the *dgk*A*-ept*A1 operon.

**Fig. 5.**
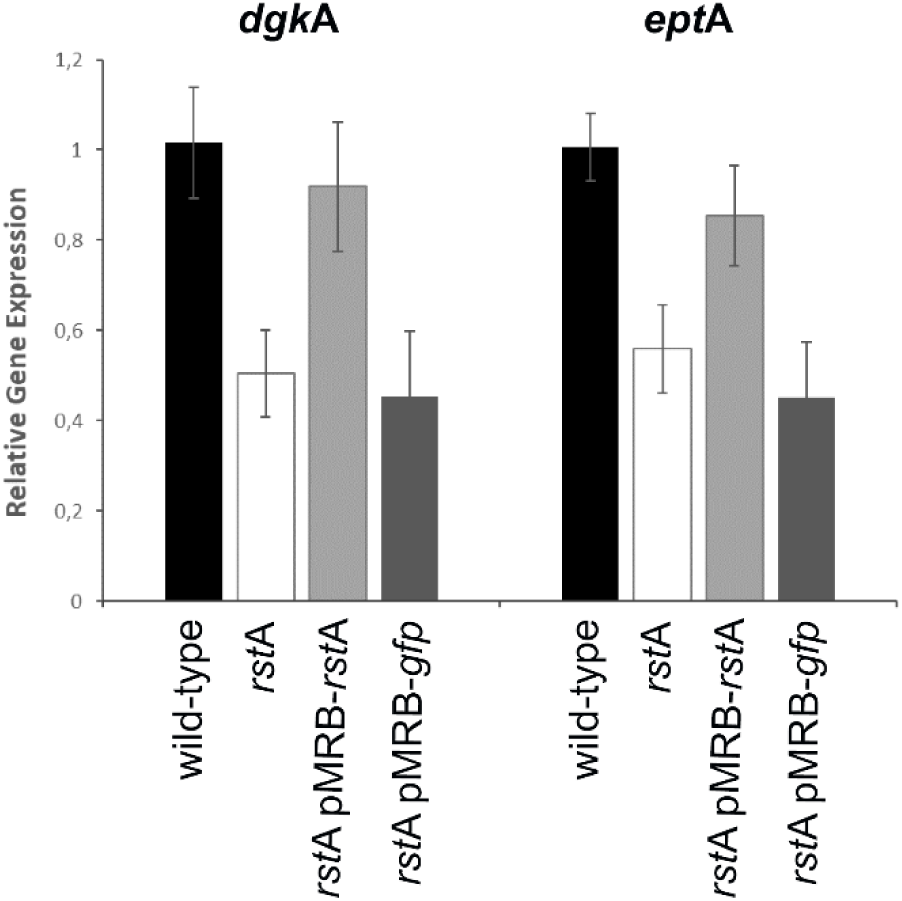
*dgk*A and *ept*A gene expression is controlled by *rst*A. Expression of *dgk*A and *ept*A genes was quantified in wild-type *V. harveyi* Th15_F5-F11 and its *rst*A isogenic mutant. The *rst*A-deletion mutant and the mutant complemented with a pMRB-gfp plasmid showed a significant decrease in expression for *dgk*A (estimate= -0.511 ± 0.173, t = -2.963, p = 0.021 and estimate = -0.561 ± 0.193, t = -2.910, p = 0.023) and *ept*A (estimate = -0.446 ± 0.136, t = -3.271, p = 0.014 and estimate = -0.553 ± 0.153, t = -3.623, p = 0.008), whereas complementation with *rst*A restored wild type expression levels (*dgk*A estimate = -0.0978 ± 0.172, t= -0.567, p = 0.589 and *ept*A estimate = -0.151 ± 0.136, t = -1.109, p = 0.304). Data were normalized using two housekeeping genes.

### Other mechanisms of colistin resistance were found in *Vibrio*

While the *rst*A*-rst*B*-glycine zipper-dgk*A*-ept*A gene cluster was conserved in 6 of the 9 sequenced colistin-resistant isolates from Thau (66,6%), three resistant isolates from our European sampling lacked both the *ept*A and *mcr* genes (*Vibrio sp.* TH21_20_OG1, TH21_20_OH4 and TH21_20A_OC7). In these strains, we detected a well-known colistin resistance mechanism, encoded by *arn*BCADTEF (transfer of L-arabinose onto Lipid A), along with the *pho*P/Q two-component system known to regulate the expression of *arn*ABCDEFT in a broad number of bacterial species ^12^ (Fig. S5). These three isolates exhibited an average nucleotide identity (ANI) of 91% with *Vibrio variabilis* strain CAIM1454, a marine bacterium previously isolated from the cnidarian *Palythoa caribaeorum*^31^. Next to *pho*P/Q and *arn*ABCDEFT, we further detected a range of other orthologs of known colistin resistance genes, such as *pmr*A/B (= *bas*S/R), *crr*A/B, *sox*R, *tol*C, *kpn*E, *mpr*F, and *ug*D genes in the pool sequencing libraries from Thau, Sylt and Ebro^12^. While we have not functionally characterized these genes, the high prevalence of colistin resistance in the absence of *eptA*-1-2-3 suggest the presence of a substantial diversity of known and probably also novel resistance mechanisms.

## Discussion

This study reveals that *eptA* genes, which encode phosphoethanolamine transferases, are both abundant and diverse in culturable bacteria isolated from European coastal environments and are widely distributed across *Vibrionaceae*. We specifically show that ancient copies of *eptA* have evolved within distinct genomic environments specific to each *Vibrio* clade and contribute to intrinsic Col-R in *Vibrionaceae*. In addition, a number of *ept*A paralogues exhibit signatures of recent mobility within *Vibrio*, highlighting the need for increased surveillance. These results fill a gap of knowledge. Indeed, while *eptA* genes had previously been described in a number of *Vibrio* species for their role in conferring colistin resistance, little was known about their distribution in coastal environments as well as in the *Vibrionaceae* family.

In European coastal waters (France, Germany, Spain), *eptA* genes displayed high diversity, with the most abundant variants found in *Vibrio* species belonging to the Harveyi and Splendidus clades. Indeed, among *ept*A sequences, we identified four novel *ept*A gene variants referred to as *ept*A*-*1 to -4 in this study. EptA-1 to -3 were expressed by colistin-resistant strains of the Harveyi clade assigned to the species *V. alginolyticus, V. campbellii, V. diabolicus, V. harveyi, V. jasicida, V. owensii,* and *V. rotiferianus.* In contrast EptA-4 was expressed by susceptible strains of the Splendidus clade (species *V. splendidus* and *V. crassostreae*). These two *Vibrio* clades naturally colonize oysters, and several species such as *V. harveyi* and *V. crassostreae* cause infections in oysters^32,33^. A remarkable contrast was observed in the geographic distribution of *ept*A gene variants at a European scale, with the active forms (*eptA*-1 to -3) only being detected in the Mediterranean area (Ebro, Spain and Thau, France), suggesting an adaptive advantage responding to specific selection pressures in that environment. However, since these genes were carried by *Vibrio* of the Harveyi clade, which are adapted to warmer seawater temperatures, the distribution of these *ept*A variants likely reflects the geographic range of *Vibrio* species along European coasts. This does not rule out the possibility that environmental factors also select for *ept*A-mediated Col-R in the Mediterranean coastal environments. No division was observed in the geographic distribution of *ept*A*-*4 (Splendidus clade), which were found in all three European environments along with a number of unknown *mcr* genes. Our data also showed that two weeks were sufficient for oysters to capture a number of Col-R genes specific to each European environment. This finding has implications for the potential transfer of AMR across Europe by oyster transport, a common and still unregulated practice in aquaculture, which has been responsible for the spread at the European scale of oyster pathogens including *Vibrio*^34^.

In the Harveyi clade, we described a novel *ept*A genomic environment where *ept*A is co-transcribed with *dgk*A under the control of the RstAB two-component signal transduction system, whose environmental triggers remain unknown. Co-expression of the *dgk*A-*ept*A-1 operon was required for Col-R, in agreement with recent results indicating that *dgk*A is needed for the *ept*A or *mcr*-1-mediated resistance to polymyxins in *E. coli*^35^, as well as in environmental isolates carrying *mcr-3* and *mcr-*7 ^36^. The underlying mechanism involves the detoxifying effect of DgkA, a diacylglycerol kinase that recycles diacylglycerol, a dead-end metabolite of Mcr/EptA proteins, into useful precursor molecules. DgkA plays a crucial role in Col-R by preventing the toxicity of Mcr/EptA by-products from inhibiting bacterial growth (for review see ^37^). We also showed that co-expression of *dgk*A-*ept*A-1 is controlled by RstAB, which enables bacteria to detect and respond to environmental fluctuations^38–40^ and triggers adaptive responses for bacterial survival^39–41^. Until now in *Vibrio*, RstAB had been shown to control motility, adhesion, biofilm formation and haemolytic activity^42^. Whereas in *Photobacterium damselae* RstAB would not control the *ep*tA-mediated resistance to colistin^43^, the homologous TCS *car*RS controls *ept*A expression and Col-R in *V. vulnificus*^25^.

Unlike in Harveyi, the conserved *eptA*-4 copy carried by *Vibrio* of the Splendidus clade, which has evolved in a completely distinct genomic environment lacking *dgk*A, appears to have lost its ability to confer colistin resistance, as also reported for *ept*A from the classical *V. cholerae* strain O395^23^. The conservation of this gene in the Splendidus clade may indicate that EptA proteins have evolved different specificities/functions along the *Vibrio* phylogeny, although we cannot rule out the possibility that the gene was not activated under our experimental conditions.

Ancient copies of *eptA* were found widely distributed in *Vibrionaceae* suggesting they contribute significantly to intrinsic colistin resistance in these bacteria. First, from our phylogenetic studies on nearly 30.000 *Vibrionaceae* genomes, we showed that only a very limited number of species (*e.g. V. natriegens)* have lost this gene during evolution. Second, despite sequence identities as low as 43% across the *Vibrio* phylogeny, many EptA variants retain a conserved role in conferring colistin resistance, as shown here for EptA-1 from the Harveyi clade in the EUCAST conditions, and reported elsewhere for species of the *Vibrio* clades Cholerae, Vulnificus, and Fisheri ^23, 25^,^26^. This gives some hints about the functional role and selective pressures encountered by this gene in nature. In the marine environment, various microorganisms, including bacteria produce cationic lipopeptides. Thus, species of *Pseudoalteromonas* are members of the oyster microbiota and the lipopeptides they produce (called alterins) are structurally and functionally similar to polymyxins^44^. Moreover, marine animals such as oyster^45^ and squid^26^ produce cationic antimicrobial (lipo)peptides and proteins as a mechanism of immune defense and to control their microbiota. Like polymyxins and alterins, some of them target the lipopolysaccharide of Gram-negative bacteria^46^. Not surprisingly, Lipid A modifications have evolved as defense mechanisms against cationic antimicrobial peptides, both in pathogens and commensals to circumvent the antimicrobial response of their animal host and competing members of the host microbiome^47^. The importance of *ept*A-mediated colistin resistance in the success of host colonization was clearly demonstrated in the squid symbiont *V. fisheri*^26^. An almost identical gene (>98% sequence identity) was sampled two times here in oysters from Thau and Ebro. It is likely that bacterial species that live in close association with marine animals like oysters, such as *Vibrio* clade Harveyi^32^, have also developed such resistance mechanisms. We also demonstrated here that conserved copies of *eptA* have evolved within *Vibrio* species in specific genomic environments, further supporting the hypothesis of an ancient acquisition of the *eptA* gene in *Vibrio* phylogenetic history and little to no interspecific horizontal gene transfer since the differentiation of *Vibrio* species. The diversity of genomic environments observed for *eptA* variants is specific to *Vibrio* clades. As we showed here for *V. harveyi*, these genomic contexts are key determinants of *EptA* expression and activity. The presence in many *ept*A genomic contexts of clade-specific two-component regulatory systems suggests that *eptA* expression responds to distinct environmental triggers in different *Vibrio* clades.

Importantly, we also evidenced recent mobility events for certain *eptA* paralogues. Indeed, beyond the ancient *ept*A copy inherited by the majority of *Vibrionaceae* species, we found that a number of species from the clades Harveyi, Cholerae, Anguillarum and Fluvialis harbor mobile *ept*A paralogues. Their genomic environment differs from the clade-specific ancient *ept*A copy, indicating they are likely expressed in response to distinct environmental signals. Although representing less than 1% of the *ept*A genes identified in *Vibrio*, the mobile *ept*A paralogues are surrounded by transposases and integrases and show signs of recent mobility both at an intra-clade and at an inter-clade level. Remarkably they co-occur with a *dkg*A gene in their direct neighborhood, suggesting functionality. This deserves particular attention since such mobile genetic elements significantly increase the risk of environmental capture, as this happened for the *mcr-*1 gene carried by a plasmid and now circulating in human pathogens.

## Conclusion

Our exploration of the genetic basis of Col-R focused on *mcr/ept*A genes, which have a putative origin in aquatic environments and currently raise concerns in a clinical context worldwide^48,49^. We found a high diversity of *ept*A variants, but only a limited number of *mcr* variants in bacteria isolated from European coastal waters. An ancient *ept*A gene copy was highly prevalent in *Vibrio* supporting a key role in *Vibrio* intrinsic resistance to colistin including strains from species responsible for most human pathologies (*V. cholerae*, *V. parahaemolyticus*, *V. vulnificus*, *V. alginolyticus*). Most often, these species harbor a *dgk*A*-ept*A operon under the control of a two-component regulatory system (RstAB in the Harveyi clade or its homolog CarRS in the Vulnificus clade). In European coastal environments, the newly described Harveyi *ept*A genomic environment was found in Mediterranean strains of *V. alginolyticus and V. harveyi,* which thrive in warm seawaters. The chromosomal location of *ept*A, the absence of mobile genetic elements at the vicinity of the *dgk*A*-ept*A genes and the evidence that the *ept*A gene polymorphism has evolved with the *Vibrio* lineage argue in favor of a low risk of horizontal gene transfer from *Vibrio* to other bacterial genera. However, the identification of mobile *ept*A paralogues in *Vibrio* genomes should warn us on a risk of mobilization outside *Vibrionaceae*. Further studies will be needed to characterize the risk associated to other Col-R mechanisms present in oyster associated bacteria. In the context of global warming, where *Vibrio* are already causing an increasing number of human disease cases in Europe^50^, we expect colistin-resistant *Vibrio* species of the Harveyi clade to proliferate. A correct understanding of the spread of Col-R and of ecological factors interfering with it in coastal areas, and of its persistence and selection in the oyster pathobiome is fundamental for the preservation of oyster farming as sustainable aquaculture in oceans exposed to global warming ^20^.

## Methods

### Animals

Specific-pathogen free (SPF) diploid oysters were produced during Spring 2021 from 166 wild genitors at the Ifremer hatchery of Argenton (PHYTNESS Ifremer research unit, France), as described in Petton (2011, 2015). Oysters were transferred after 6 weeks to the Ifremer nursery of Bouin (EMMA Ifremer research unit, France), where they were maintained under controlled biosecured conditions with filtered and UV-treated seawater enriched in phytoplankton (*Skeletonema costatum, Isochrysis galbana,* and *Tetraselmis suecica*). Before transfer to the field, the SPF status of the animals was confirmed by (i) the absence of OsHV-1 DNA detection by qPCR, (ii) a low *Vibrio* load (∼10 CFU/mg of oyster tissue) determined by isolation on selective culture medium (TCBS agar). The absence of World Organisation for Animal Health (WOAH) listed parasites (*Bonamia* sp., *Marteilia* sp., *Perkinsus* sp., *Mikrocytos* sp. and *Haplosporidium* sp.) was confirmed on histological sections of 114 oysters by an independent laboratory (LABOCEA, France). Oysters were observed to remain free of any abnormal mortality until use.

### Study Area, Sample Collection and Microbial Isolation

The same SPF oyster batches (*i.e.* oysters with identical life history traits) were deployed at the juvenile stage (6 months old) in three different European locations used for oyster culture: SYLT in Germany (N 55° 1’ 42.539’, E 8° 26’ 1.953’), THAU lagoon (N 43°26.058’, E 003°39.878’) in France, and EBRO delta (N 40°37.106880’, E: 0°37.318320’) in Spain. Two to three weeks exposures in the environment were performed at the end of the year 2021 (THAU: Bouzigues, from 4/10/2021 to 18/10/2021, EBRO: Alfacs bay, from 8/10/2021 to 25/10/2021, and SYLT: Königshafen, from 22/12/2021 to 10/01/2022). No oyster mortalities were recorded during the field exposures. After being exposed to their respective environments, 60 oysters were collected at each sampling site. Tissues were homogenized in artificial seawater (ASW: 400 mM NaCl, 20 mM KCl, 5 mM MgSO4, 2 mM CaCl2) using an ultra-Turrax apparatus^51^. The homogenates were pooled and plated in replicates on TCBS agar (Difco™) agar plates for *Vibrio* isolation and on marine agar (Difco™) as a non-selective agar for marine bacteria. Duplicate plates were incubated at 20°C or 37°C. Bacteria were isolated after 24-48h (Table S1). After colony purification on Zobell medium (ASW supplemented with 0.4% bactopeptone and 10% yeast extract, pH 7.8), a glycerol stock of each isolate was conserved at -80°C. Up to 48 isolates were conserved per condition of isolation (TCBS 20°C, marine agar 20°C, TCBS 37°C, marine agar 37°C).

### DNA extraction, pool-sequencing, and whole genome sequencing

Bacterial colonies isolated from oyster flesh were cultured overnight in liquid Zobell medium at either 20°C or 37°C, according to their conditions of isolation. For pool-sequencing, bacterial cultures were pooled in equal amounts according to geographic site, culture medium and temperature of isolation. DNA was extracted with the NucleoSpin Tissue kit for DNA from cells and tissue (Macherey-Nagel). For single genome sequencing, DNA was extracted from cultures with the MagAttract HMW DNA Kit (QIAGEN, France). The extracted DNA was quantified using a Qubit High Sensitivity Assay Kit (Life Technologies, Carlsbad, USA), and sequencing was carried out at the Bio-Environment platform (University of Perpignan Via Domitia) using the Nextera XT DNA Library Prep Kit (Illumina) according to the manufacturer’s instructions, with 1 ng of DNA. The quality of the libraries was checked using a High Sensitivity DNA chip (Agilent) on a Bioanalyzer. Sequencing was performed on a NextSeq 550 instrument (Illumina) in 2x150 paired-end mode, resulting in an average mean reads of 45,000,000 bp for pool sequencing, and 243 Mb for whole genomes (mean coverage 47X).

### Bioinformatics Analysis tools

#### Pool-sequencing

FastQC was used to check the quality of reads, followed by trimming using Trimmomatic V-0.38^52^ to trim leading/trailing bases with quality scores below 30. The recovered reads were assembled into contigs using MEGAHIT V-1.2.9^53^. The Meta-marc tool (database model 3) was then used to identify the contigs carrying the *eptA/mcr* resistance genes (Fig. S6). The contigs recovered carrying the *eptA/mcr* resistance genes were annotated using Prokka and predicted coding sequences was specifically re-analyzed for colistin resistance genes using Meta-marc. Sequences annotated as *ept*A and *mcr* according to Meta-marc and Prokka, were validated by BlastP against the NCBI and the CARD database V-3.2.7. Subsequently, we eliminated incomplete *ept*A and *mcr* sequences (*i.e.* partial sequences missing a 5’ and/or 3’ region) from our analysis. Full length gene sequences were translated *in silico* and the resulting amino acid sequences were aligned by MAFFT V-7.407 (https://ngphylogeny.fr/tools/). A phylogenetic tree was generated through maximum likelihood analysis of deduced EptA/Mcr amino acid sequences using PhyML V-3.0 (https://ngphylogeny.fr/tools/) with the WAG model and 100 bootstrap replicates. Pool-sequencing raw data and complete *eptA* gene sequences were deposited on GenBank under accession numbers SAMN37810832 to SAMN37810840 and OR578979 to OR579029, respectively.

#### Whole genome sequencing of single bacterial isolates

The quality assessment and reads trimming steps were performed as described for pool-sequencing. The obtained reads were then assembled into contigs using Spades V-3.15.4 within the Galaxy Europe platform^54^. Default parameters were used (’Isolate’, ’Automatic k-mer selection, Phred quality offset adjustment, and coverage cutoff in the assembly of individual bacterial genomes). In order to detect the presence of the *ept*A or *mcr* genes, the assembled genomes were annotated in MAGe (https://mage.genoscope.cns.fr/). When feasible, a taxonomic affiliation was assigned to the selected isolates at the species level using Average Nucleotide Identity (ANI) and DNA:DNA hybridization (dDDH) percentages. This analysis was conducted utilizing Defast (DDBJ Fast Annotation and Submission Tool) available at https://dfast.ddbj.nig.ac.jp/, in conjunction with genome clustering tools through MAGe and the reference strain genomes obtained from Type Strain Genome Server (TYGS) (https://tygs.dsmz.de/). Raw reads and genome assemblies were deposited on ENA under project number PRJEB67316.

#### Genetic diversity of detected *ept*A and *mcr* genes, and their genomic environment

In order to investigate the diversity of the detected *ept*A/*mcr* sequences, we conducted a comparative analysis of the amino acid sequences encoded by genes found in the assembled pool-sequencing data. First to determine a sequence identity cut-off enabling the discrimination of Mcr variants, we used a total of 177 *mcr* sequences sourced from NCBI and CARD databases whose deduced amino acid sequences showed homology with Mcr-1-10 (Table S3). Next, Mcr/EptA sequences were searched using diamond blastp (v2.1.9.163)^55^ against a collection 29244 *Vibrionaceae* assemblies (GenBank July 2023) using 30% identity and 50% coverage as selection criteria. Multiple proteins alignment was then performed using FAMSA (v2.2.2)^56^ and trimmed using trimAI with gappyout method (v1.5)^57^. A phylogenetic tree was constructed using fasttree with LG model (v2.1.11)^58^ and visualization was done using iTOL^59^.

Additionally, we examined the genomic environment of all *ept*A/*mcr* genes from the 29244 *Vibrionaceae* assemblies. Neighbor genes from +4 to -4 were clustered with the diamond cluster module. After filtering out incomplete environments, i.e. those which lacked genes at +4 or -4 due to contig breaks, the different genomic environments were positioned on a *Vibrionaceae* phylogenetic tree. For that, a multilocus sequence analysis of *Vibrionaceae* genomes was performed based on 8 genes recommended by *Vibrio* Clade 3.0^60^, namely *fts*Z, *gap*A, *mre*B, *rpo*A, *top*A, *gyr*B, *pyr*H, *rec*A. Alignment of the 8 concatenated nucleic sequences was done with halign (v3.0.0)^61^. A phylogenetic tree was constructed with fastree with a GTR model. iTOL was used for visualization. ISfinder (https://isfinder.biotoul.fr/blast.php), PlasmidHunter^62^ and MetaPhinder-2.1 (https://cge.food.dtu.dk/services/MetaPhinder/) were used to search the insertion sequences, mobile elements and bacteriophage sequences within a 25,000 bp average region surrounding the *ept*A gene.

#### Data normalization and statistical analysis

Bowtie2 V-2.5.1^63^ was employed with its default settings to align reads to the gene sequences of interest. This approach provides read counts for each identified sequence (Table S4). To normalize the number of reads of each identified sequence, the following formula was used:

Normalized data= (mapped reads of each sequence / total assembly reads) x 10^6609^

Furthermore, we investigated the geographical structuring of molecular diversity in EptA and Mcr sequences by analyzing the distance matrix underlying the amino acid sequences phylogeny (see above). This involved conducting pairwise comparisons of distances within and between sites using a permutational analysis of variance (PERMANOVA). The PERMANOVA was implemented in the R package *vegan*, utilizing the *adonis*2 function. (https://github.com/vegandevs/vegan). Initially, we conducted an analysis of variation using all sites collectively. Subsequently, we performed pairwise comparisons to detect low-level structuring.

### Colistin susceptibility testing

MICs of colistin were determined against *Vibrio* isolates and recombinant *E. coli* strains constructed in the present study following the microdilution assay from the EUCAST V14.0 guidelines. Colistin sulfate (Thermo Fisher) corrected for activity units was tested in the range of 0.125 to 16 µg/ml, at 35°C, in cation-adjusted MHCA. Each well was seeded with 5.10^5^ CFU/ml. MIC values are expressed as the lowest colistin concentration tested that causes 100% of growth inhibition after a 18h incubation. In parallel, we performed MIC determination in Zobell medium, which mimics the seawater composition. For quality control (QC), we used *Escherichia coli* O6 (ATCC 25922) with the colistin QC MIC range provided by EUCAST v14.0 (0.25-1 µg/ml). In the absence of colistin breakpoint for *Vibrio*, we used EUCAST breakpoints for *Enterobacterales* resistance in tables V14.0 (MIC > 2 µg/ml).

For a rapid screening of Col-R on 139 bacterial isolates from the Thau lagoon, France, 184 from Ebro, Spain and 96 from Sylt, Germany and control *Vibrio* strains from previous studies (see Tables S5-S6), we used the same liquid broth inhibition assay with fixed colistin concentrations (5 µg/ml). Screening was performed in Zobell medium at 20°C or 37°C, according to conditions of isolation. Assays were performed in duplicate wells. Strains were considered resistant when duplicate wells grew in the presence of 5 µg/ml colistin (*i.e.* 2.5 fold the clinical breakpoint) after a 18h incubation.

### Cloning *dgk*A and *ept*A genes

Gene variant *ept*A-1 and the operon *dgk*A-*ept*A-1 were amplified by PCR from colistin resistance strain *Vibrio owensii* Th15_Z_G08 (Table S5). The *ept*A-4 variant was amplified from *Vibrio splendidus* 7T7_2. Primer sets used for gene amplification are listed in Table S6. PCR were performed in a 25 µl total volume under the following conditions: initial denaturation at 95°C for 30 s, followed by 35 cycles of 95°C for 5 s, 69°C for 30 s, and 72°C for 30 s and by a final extension at 72°C for 2 min. Amplicons were cloned in the pBAD-TOPO expression vector (Invitrogen, France) under the control of the pBAD inducible promoter and transformed into *E. coli* TOP10 competent cells according to manufacturer’s instructions. Recombinant colonies were tested for the presence of specific *ept*A and *dgk*A genes by standard PCR and by whole plasmid sequencing using Oxford Nanopore Technologies sequencers (Eurofins, France).

### Heterologous expression of *dgk*A and *ept*A genes in *E. coli*

Recombinant *E. coli* TOP10 carrying *ept*A-1, *dgkA*-*ept*A-1 or *ept*A-4 in the pBAD-TOPO vector were tested for colistin resistance. Basically, bacterial cells were cultured at 37°C in Luria-Bertani (LB) broth in the presence of 2 % arabinose to induce the pBAD promoter. Recombinant bacteria were considered colistin-resistant if they were able to grow in LB containing 5 µg/ml colistin in the microtiter plate assay described above.

### Generation of mutants in the regulation systems *rst*A/B

The *Vibrio* strain TH15_F5_F11 carrying *dgk*A-*ept*A-1 preceded by *rst*A/B (Col-R) was grown at 37°C in LB or LB-agar (LBA) + 0.5 M NaCl. *E. coli* strains were grown at 37°C in LB broth and on LB medium for cloning and conjugation experiments. Chloramphenicol (Cm, at 5 or 25 μg/ml for *Vibrio* and *E. coli,* respectively), thymidine (0.3 mM) and diaminopimelate (0.3 mM) were added as supplements when necessary. Induction of the P_BAD_ promoter was achieved by the addition of 0.2% L-arabinose to the growth medium and, conversely, was repressed by the addition of 1% D-glucose where indicated. All plasmids used or constructed in the present study are described in Table S5. Gene deletion was performed by allelic exchange using the pSW7848T suicide plasmid^64,65^. To this end, two ≈500 bp fragments flanking the target gene were amplified (Table S6), cloned into pSW7848T as previously described^66^, and transferred by conjugation from *E. coli* as donor to *Vibrio* as recipient. Subsequently, the first and second recombination’s leading to pSW7848T integration and elimination were selected on Cm/glucose and arabinose-containing media, respectively. Deletion mutants were screened by PCR using external primers flanking the target gene. For the complementation experiments, the gene was cloned into the stable pMRB plasmid, resulting in constitutive expression from a P_LAC_ promoter^67^. Conjugations between *E*. *coli* and *Vibrio* were performed at 37°C^64^.

### RT-qPCR

In this study, we employed the DirectZol RNA Miniprep kit (R2051) provided by ZymoResearch to extract total RNA from Trizol conserved samples obtained from both wild and mutant strains, following the manufacturer’s instructions. The extraction process was performed in duplicate for each condition at two different growth stages, specifically the exponential and stationary phases. To eliminate any genomic DNA contamination, the RNA was treated with DNase I. To determine the concentration of the total RNA, we used a NanoDrop spectrophotometer from ThermoFisher Scientific. The cDNA was produced using M-MLV Reverse Transcriptase M1302 (Sigma-Aldrich, France) with 1 µg of extracted RNA. Real-time quantitative PCR (qPCR) was performed at the MGX platform in Montpellier. The MGX platform employed the Light-Cycler 480 System from Roche. The primers used for qPCR are listed in Supporting Information (Table S6). To analyze the relative expression levels, we employed the 2^-ΔΔCq^ method developed by Pfaffl in 2001^68^. For normalization, two genes (6PFK (VS_2913) and CcmC (VS_0852)) were chosen due to their constitutive expression across various conditions in both the RNAseq and qRT-PCR analyses^69^. For our own study, we designed and validated specific primers for these genes on *Vibrio harveyi* Th15_F5_F11 (Table S6).

## Acknowledgements

This research was supported by the ERA-net cofund Aquatic Pollutants program of the combined European Joint Program Initiatives JPI Oceans, JPI Water and JPI AMR. It has received funding from the European Union’s Horizon 2020 research and innovation programme under grant agreement No. 869178-AquaticPollutants. This study is part of the « Laboratoire d’Excellence (LabEx) » TULIP (ANR-10-LABX-41) framework.

We are grateful to Jean-François Allienne, Margot Doberva and Michèle Laudié from the Bio-Environment platform (UPVD, Région Occitanie, CPER 2007-2013 Technoviv, CPER 2015-2020 Technoviv2) for technical support in library preparation and sequencing, as well as Oriane Chevalier and Marc Leroy (IHPE) for crucial technical help. We thank Prof. Frédérique Le Roux (Université de Montréal, Canada), Prof. Carmen Amaro (Univ. de Valencia, Spain) and Dr. Dominique Hervio (Ifremer, Brest, France) for kindly providing *Vibrio* strains. We thank the qPHD platform/Montpellier genomix for access to qPCR. We thank the Ifremer’s hatchery team EMMA (PMMLT, PMMB) of La Tremblade and Bouin for the production of Pacific oysters. We finally thank the Regional Committee of Mediterranean Shellfish Aquaculture (CRCM) and the Ifremer for access to the shellfish tables and for boats.

## Author contributions

Conceptualization: S.J., B.V., T.M.A., M.C., W.K.M., D.D.G.

Methodology: S.J., B.V., B.C., A.B.K., G.D., A.M., P.B., L.Y, T.P., F.D., A.G., V.L., C.G., D.C.A., K.H., L.E., C.G., R.O., P.J., E.J.M., M.D., C.G., T.E., T.M.A., M.C., W.K.M., D.D.G.

Investigation: S.J., B.V., B.C., A.B.K., G.D., A.M., P.B., L.Y, T.P., F.D., A.G., V.L., C.G., D.C.A., K.H., L.E., C.G., R.O., P.J., E.J.M., C.G., T.E., T.M.A., M.C., W.K.M., D.D.G.

Visualization: S.J., B.V., B.C., A.B.K., G.D., A.M., P.B., L.Y, T.P., F.D., A.G., V.L., C.G., D.C.A., K.H., L.E., C.G., R.O., P.J., E.J.M., C.G., T.E., T.M.A., M.C., W.K.M., D.D.G.

Funding acquisition: W.K.M., D.D.G., C.G., V.L., A.B.K. F.D. Administration: A.B.K., V.L, W.K.M., D.D.G.

Supervision: A.B.K., W.K.M., C.G., D.D.G.

Writing– original draft: S.J., D.D.G.

Writing – review & editing: S.J., B.V., B.C., A.B.K., G.D., A.M., P.B., L.Y, T.P., F.D., A.G., V.L., C.G., D.C.A., K.H., L.E., C.G., R.O., P.J., E.J.M., M.D., C.G., T.E., T.M.A., M.C., W.K.M., D. D.G.

## Ethics approval

The animal (oyster *Crassostrea gigas*) testing followed all regulations concerning animal experimentation. The authors declare that the use of genetic resources fulfill the French regulatory control of access and EU regulations on the Nagoya Protocol on Access and Benefit-Sharing (TREL2302365S/750, ABSCH-IRCC-FR-266230-1).

## Data availability

Targeted gene sequences (*eptA*) and pool-sequencing raw data were deposited at GenBank under accession numbers OR578979 to OR579029 and SAMN37810832 to SAMN37810840, respectively. Genome raw data and assemblies were deposited at the European Nucleotide Archive (ENA) under project accession no. PRJEB67316 (ERR12116510 to ERR12116518) and are available on MicroScope plateforme MaGe (“Magnifying Genomes”) https://mage.genoscope.cns.fr/.

## Code availability

Software’s and databases used in this paper are openly available in Galaxy Open platform hosted by our institute https://galaxy-datarmor.ifremer.fr/. Specific source codes for antimicrobial resistance gene identification are also available on GitHub https://github.com/lakinsm/meta-marc.

