## Supplementary data for "*Vibrio* are a potential source of novel colistin-resistance genes in European coastal environments"

**Table of content**

Fig. S1 The pourcentage of Col-R bacterial isolates across European oyster farms. p 3

Fig. S2. Phylogeny of EptA/Mcr sequences based on medium culture and genus identification. p 4

Fig. S3. Multiple sequence alignment of EptA-1 to -4 with EptA orthologs in *Vibrionaceae*. p 5

Fig. S4. Percentage of EptA paralogues in species of *Vibrionaceae*. p 7

Fig. S5. Alternative colistin resistance mechanisms identified in *Vibrio* strains TH21_20_OC7, TH21_20_OH4, and TH21_20A_OG1. p 8

Fig. S6. Bioinformatics analyses for identifying *ept*A and *mcr* gene sequences in Poolseq data. p 9

Table S1. Metadata on *mcr/eptA* unique sequences found in Pool sequencing data p 10

Table S2. Genbank ID of nucleotide sequences of *mcr*/*ept*A genes p 11

Table S3. List of *mcr*/*ept*A unique sequences found in pool sequencing data p 13

Table S4. Amino acid sequence identity between EptA variants. p 15

Table S5. Strains and Plasmids p 16

Table S6. List of primers used in this study p 19

A

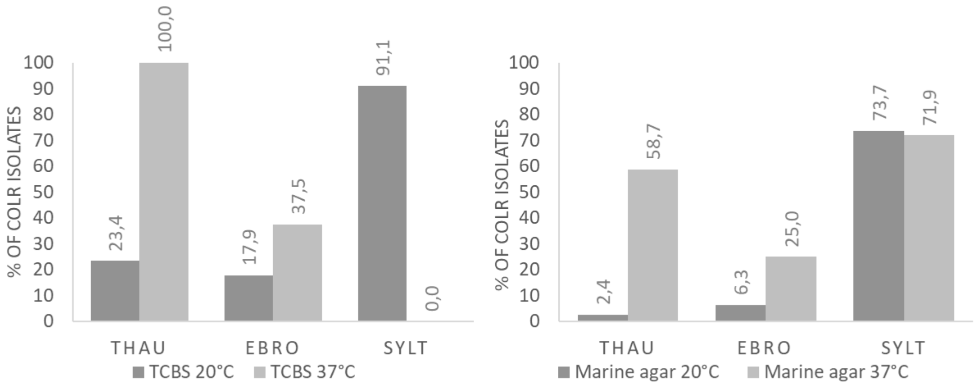


B

| | Estimate | Std. Error | z value | Pr(>\|z\|) |
|---|---|---|---|---|
| **Intercept** | -4.0118 | 1.0429 | -3.847 | 0.00012 |
| **Temp37** | **4.3876** | **1.0661** | **4.115** | **3.86e-05** |
| **MediumTCBS** | **2.8581** | **1.0727** | **2.664** | **0.00771** |
| **SiteEBRO** | 1.3510 | 1.0855 | 1.245 | 0.21328 |
| **SiteSYLT** | **5.0414** | **1.1658** | **4.324** | **1.53e-05** |
| **Temp37:MediumTCBS** | -0.3747 | 0.8405 | -0.446 | 0.65571 |
| **Temp37:SiteEBRO** | **-2.8585** | **1.1307** | **-2.528** | **0.01147** |
| **Temp37:SiteSYLT** | **-4.4789** | **1.2501** | **-3.583** | **0.00034** |
| **MediumTCBS:SiteEBRO** | -1.7907 | 1.1590 | -1.545 | 0.12233 |
| **MediumTCBS:SiteSYLT** | -1.5604 | 1.3025 | -1.198 | 0.23090 |

**Fig. S1 The percentage of Col-R bacterial isolates across European oyster farms.**

(A) % Col-R isolates on TCBS (left panel) or marine agar (right panel). (B) Results of a binomial general linear model showing the effect of environmental origin (Site), temperature (temp) and culture medium (Medium) plus their two-way interactions on the proportion of Col-R bacteria.

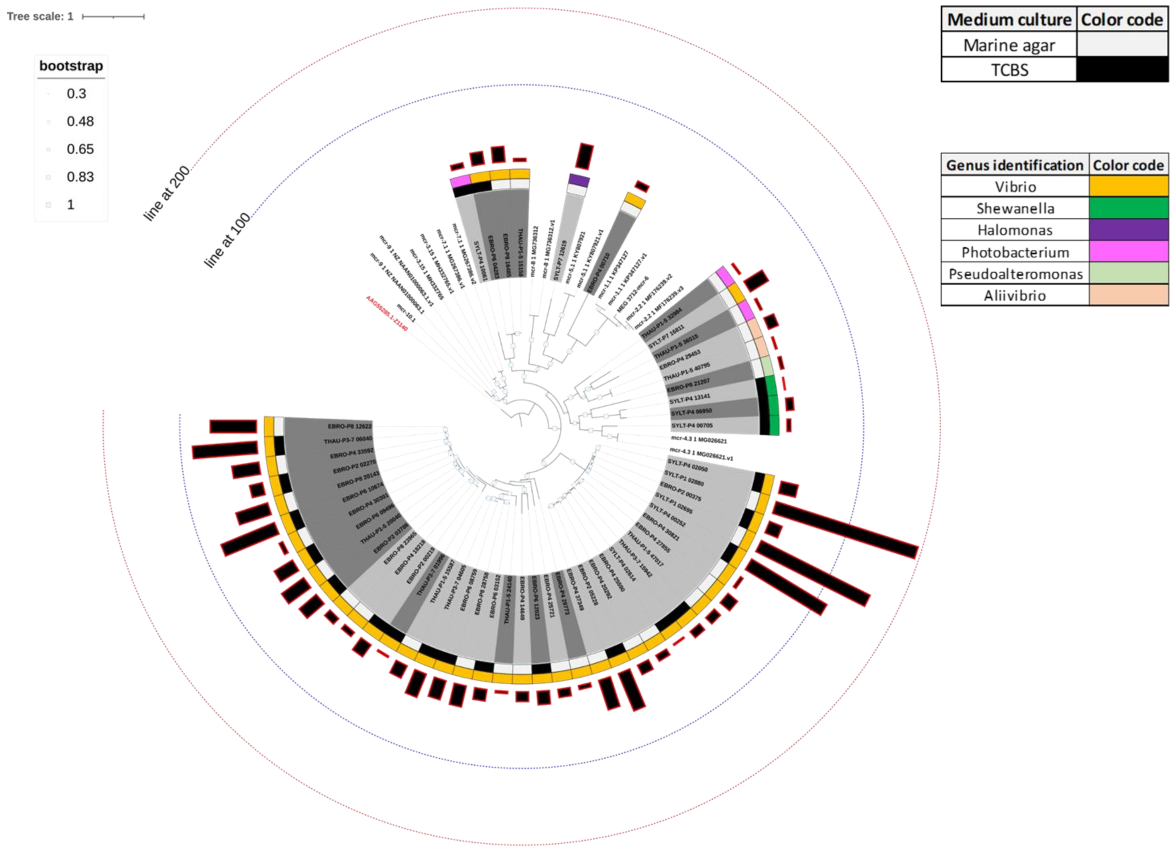


**Fig. S2. Phylogeny of EptA/Mcr sequences based on medium culture and genus identification.**

This figure represents conditions of isolation (back and grey boxes), genus identification (color boxes) and read counts (black bars with a red outline). The phylogenetic tree was generated through maximum likelihood analysis of deduced EptA/Mcr amino acid sequences using PhyML V-3.0 with the WAG model and 100 bootstrap runs. Amino acid sequences were extracted from the annotated file generated from pooled sequencing of bacteria isolated from both Marine agar and TCBS medium across three European regions. This figure represents conditions of isolation, genus identification and read counts. Tree was visualized using the iTOL online website. Blue line at 100 and red line at 200 represent the numerical scale of normalization.

```
EptA-1                  -------MKTIELPTKGLSYVTFTLLLALYFALVVNIPIYKELVHILTNLDQVKIGFIIT   53
EptA-2                  -------MKNITFPYKGISYVTFTLFLALYFALVVNIPIYKELSHILSNLDEVKIGFIIT   53
EptA-3                  -------MKTHNLQQKGISYVTFTFLLALYFALVVNIPIYKELNGIFSSLDNVKIGFIIT   53
EptA-4                  ----------MNLPSLNMSINKLPFVLAVYYLLVINIPLSLELFSIVQASKSESVAFLIS   50
A. fischeri             MLVLSTLISKIKNHKISMNMVSLITIVSLYFAVAFNYPINEKIYQLSQ----GQEIFLFL   56
V. cholerae El Tor      ------MVIMPTFNIKPFSYLQLVFLLAAYFALPLNLPVYAQLAHIFNQSTSLDWGFALS   54
V. vulnificus           ------MNRIKTLFNRECSYTTFTLVLAIYFATIVNLPIYKELFSIISHLDSVKVGFVIS   54
V. parahaemolyticus     -------MKTVDSPKKGIPYVAFTVLLALYFALVVNIPIYKELIGIFSKLDEVKIGFIIT   53
                                             :  .:: *:   .* *:  ::  :       .  * :

EptA-1                  IPIFFFAALNFLFNLFSWPWISKPFFILLLITSGIVSYASYNYGTLFDTDMITNIVETDT   113
EptA-2                  IPIFFFAALNFLFNLFSWPWLSKPFFIVLLVVSSMVSYASYNYGTLFDTDMITNIVETDT   113
EptA-3                  IPIFFLAALNFLFNLFSWPWVGKPFFIILLIISSMVSYASYNYGTLFDSGMIANIAETDS   113
EptA-4                  IPIFFLAAFNFIFQVFNWPIFSKPFFIFLLITSTLVSYSMFNYGIYVDYGMIENVFETNN   110
A. fischeri             TPALLTCAFIIIFSFIAFPYVFKGIVVVLTITSAMAFYAALQYNTMFDYAMIENIFETNV   116
V. cholerae El Tor      IPLFFLFALNFIFQIFSWPYLFKPFFAVLLVLSALISFAGFQYGVIVDQDMLVNVLETDR   114
V. vulnificus           VPIFFLAALNFLFNLFSWPWLSKPFFAVLLLLSAMVSYAGYNYGTLFDYGMIANIMETDS   114
V. parahaemolyticus     IPIFFFAALNFLFNLFSWPWVGKPFFILLLVVSSLVSYAGYNYGTLFDSGMIANIVETDS   113
                         * ::  *: ::*..: :* . * :. .* : * :  ::  :*.  .*  *: *: **:

EptA-1                  SEASSYFSAYSLIWTVLMGVLPALLVFKVKFQSLKGNWLRFTLTKIVSMLASLVIIAVIA   173
EptA-2                  SEASSYLSAYSVIWTLLMGMIPAALVFKVKLQPLNGQWLRFVLTKVVSMLASLAVIAVIA   173
EptA-3                  SEASSYLSTYSVIWTLLMGGVPAMIVYKVKLQSQRHQLLYFSLTKLVSMLSSLAVIAVIA   173
EptA-4                  GEAASYVSTHSLLWLLVMGGIPSLILLLTKLK--RESWKDFFVWKSIGLLSSLIVIAIIA   168
A. fischeri             DEASSYLSSASIGYLIVFGLIPSILLLKVNIVR-HASWLKELFHRAILMSVAIVGLLLIS   175
V. cholerae El Tor      GEAGSYLTIYSVLWLLGFGLVPALALLFTPIRPE-KSALRFLMKKGLSMLVSVVVIGVIA   173
V. vulnificus           SEAGSYLSAYSVIWTLLMGVVPALLVCKLNLS-LSGSTLHMVAKKAASMLASLAVIAVIA   173
V. parahaemolyticus     SEASSYLSTYSVVWATLMGVIPALIVFKVKLQPQRGQWLRFVLTKLVAMLASLAVIAVIA   173
                        .**.**.:  *: :   :* :*:  :    :     .       :   :  ::  : :*:

EptA-1                  GLYYQDYASVGRNNSYLKKIIIPTQYVYAISSYVKETYLTTPQPYREIGTDAKQSQLALE   233
EptA-2                  GLYYQDYASVGRNNSYLKKVIIPTQYVYSISSYVKAKYLTTPEPYREIGLDAKQSSTALQ   233
EptA-3                  SVYYQDYASVGRNNSYLKKIIIPTQFVYSATGYVKEKYFTTPEPYREIGIDAKQSDSAVK   233
EptA-4                  GLFYKDYVSIGRNNSHIKKMIIPTEYVASTVKYINNTYVKEPIPYQELGLDAQLKPEAK-   227
A. fischeri             VFYFKDYASIGRNNSYLNKMINPAD-AFNTVKYIKNEYLTAPLEYITIGEDAVVTPA---   231
V. cholerae El Tor      GLYYQNYSSVGRNNSSLKKMIIPTHFLYSSFGLVKQRYFTEPMVYQEIGQDAQQKPSAVR   233
V. vulnificus           GLYYQDYASIGRNNTHLKKMIIPTQYVYSATKYVRDTYFSTPQPYRQLGLDAKQSELALQ   233
V. parahaemolyticus     GLYYQDYASVGRNNSYLKKMIIPTQYVYSATSYVKENYLTTPQPYREIGTDAQQSSTALQ   233
                         .::::* *:****: ::*:* *:.        :.  *.. *  *  :* **  .

EptA-1                  QAKEKPTLVVFVLGETARTQNYQINGYDRETNPYTSKLDVISFQDVSSCGTATAVSVPCM   293
EptA-2                  QANGKPTLLVFVVGETARTQNYQLNGYERETNPYTSKLDVVSFKDVASCGTATAVSVPCM   293
EptA-3                  QAQNKPTLLVFVVGETARTQNYQLNGYKRETNPYTSQLDVISFQEVASCGTATAVSVPCM   293
EptA-4                  -QAEKPTLLVFVLGETARVYNYQYYGYEKETNAHTKPYNPIFFSDVQSCGTATAVSVPCM   286
A. fischeri             -KNGKPTLMVILVGETARAQNSAYNGYERETNPYTKDLGLITFQNVTSCGTATAHSLPCM   290
V. cholerae El Tor      QATQKPTLVFFVLGETARVQNYQYLGYPRDTNAYTAPFQPIFFKDVASCGTATAVSVPCM   293
V. vulnificus           QADQKPTLLVFVVGETARSQNYQANGYARPTNQYTNELGMISFQDVSSCGTATAVSVPCM   293
V. parahaemolyticus     QAQDKPTLLVFVVGETARTQNYQLNGYERETNPYTSQLDVISFKDVASCGTATAVSVPCM   293
                            ****:.:::*****  *    ** : ** :*     : *.:* ******* *:***

EptA-1                  FSQLPRTDFDRSTADNQDNALDIMQRAGIDLMWKDNDGGDKGVAHKINKMDVDRSQKNEQ   353
EptA-2                  FSQLTRSDFDRSRADNQDNALDIMQRAGVDLLWKENDGGDKGVAHNITKIEVDRSQKNEL   353
EptA-3                  FSQLTRSEFERTKADNQDNALDIMQRAGVDLIWKENDGGDKEVAHKIKKIEVNRKQTSPL   353
EptA-4                  FSNMNRSNYDRDKAYNQDNVVDIMNRAGIHSIWREHDGGDKAVAHRIKEMTLVAKDSDPL   346
A. fischeri             FSNMTRTSYNKDRANNQDNALDVLTHAGVKAVWIDNDGGDKAVAKNIEKEMITDHSESNL   350
V. cholerae El Tor      FSNMNRQNFDRSRADNQDNVLDILQRAGISLLWKENDGGDKNVAKNIPLKELARDNREGI   353
V. vulnificus           FSNMTRSNFDRPTADNQDNALDILNRAGVSLLWKENDGGDKDVAKNIPLFTVDRSRKDAL   353
V. parahaemolyticus     FSQLTRNQFDRKQADNQDNALDIMQRAGIDLLWKENDGGDKEVAHKIKKIEVDRKQQNAL   353
                        **:: * .:::  * ****.:*:: :**:  :* ::***** **:.*    :     .

EptA-1                  CNGSTCYDIALLNNLDDQIASMKGNRLVALHLIGSHGPTYFQRYPKDKAFFQPDCPRADI   413
EptA-2                  CNGSTCYDMALLNNLDTQISSMKGNRVIAMHLIGSHGPTYFQRYPKDKAFFQPDCPRADI   413
EptA-3                  CNGQTCYDMALLDHFDQEVADMHGNRVVAMHLIGSHGPTYFQRYPRDKAFFKPDCPRADI   413
EptA-4                  CNNNVCYDTAMLENFEQDTQDLKQDSIIFYHISGSHGPTYFERYPDDHKKFTPDCARADI   406
A. fischeri             CNGSSCYDEILLKGLDKRIAETQGNQVYALHMIGSHGPTYWKRYPKEMAVFGPACNRSDI   410
V. cholerae El Tor      CDGDTCYDIAMLENLDQEIATQQGNRMIFMHFIGSHGPTYFKRYPKEMAVYQPDCPRADI   413
V. vulnificus           CNGSTCYDMALLENFEQDVENLKGNRVLALHLIGSHGPTYFQRYPKDKAAFMPDCPRADI   413
V. parahaemolyticus     CNGQTCYDMALLSDFDQEVSNMNGNRVVAMHLIGSHGPTYFQRYPKEKAFFQPDCPRADI   413
                        *:.. ***  :*. ::      : : :   *: *******::*** :   : * * *:**
```

```
EptA-1               ENCSVEQIVNTYDNTIRYTDFVLDQTIAKLKALEDKYNTAMIYVSDHGESLGENGLFLHG   473
EptA-2               ENCSVEQIVNTYDNTIRYTDFVLDQTIAKLKTLEDKYNTAMIYVSDHGESLGENGLFLHG   473
EptA-3               ENCSVEQIVNTYDNTIRYTDLVLAETINKLKALESQYNTALIYVSDHGESLGENGLFLHG   473
EptA-4               ENCTKEEVVNTYDNTILYTDFFLSQAIQKLEKLTDKYNVALMYISDHGESLGENGVYLHG   466
A. fischeri          ENCSDAEITNVYDNTLVYTDYVIAQTVKKLQSYSDKYNVVMLYISDHGESLGENGLYLHG   470
V. cholerae El Tor   ENCSVEQIVNTYDNTIRYSDYVMSQLLAKLDSLQDRYNTALIYISDHGESLGENGLFLHG   473
V. vulnificus        ENCSVEQIVNTYDNTILYTDYVLSQTINKLKALEERYNTALIYVSDHGESLGENGLFLHG   473
V. parahaemolyticus  ENCSVEEIVNTYDNTIRYTDFVLEQTINKLKTLEDKYNTALIYVSDHGESLGESGMFLHG   473
                     ***:  ::.*.****: *:* .: : : **.   .:**..::*:*********.*::***

EptA-1               MPYGLAPDFQKRVPMILWMSPGFKQAKHINTDCLEQEAQQTATHSHDNVFHSLLGIMDVE   533
EptA-2               MPYGLAPDFQKRVPMMLWMSPGFKQAKHINSDCLQQEAQQTATHSHDNVFHSLLGIMDIE   533
EptA-3               MPYGLAPDFQKRVPLLMWLSSGFKQVKQINTDCLSNEAQKAGQYSHDNVFHSLLGVMDVQ   533
EptA-4               MPYSLAPKEQTHVPMILWMSDGFAAQKRISETCLRKAG-KEQSFSHDNLFDSLLGLMDVQ   525
A. fischeri          APYMMAPKEQTHVPWYIWMSDDYANQKGIDKAAVIDNS-ATGDYSHDNLFHTILGLYGVT   529
V. cholerae El Tor   MPYSLAPEYQTRVPLLIWMSSGFSQSKGIDVECLRSNS--ELPYAHQNLFHSLLGVMDVS   531
V. vulnificus        MPYSLAPEYQTKVPLMLWMSPGFAQAKQINADCLKQQAKQTDTYSHDYIFHSLLGVMNVS   533
V. parahaemolyticus  MPYGLAPDFQKRVPLIMWMSPSFKQAKHINTDCLSKEAQNAGKYSHDNVFHSLLGIMDVK   533
                      ** :**. *.:**  :*:* .:   * *.  .: . .     .:*: :*.::**:  .:

EptA-1               TDAYDGDLDIFKTCRTS--       550
EptA-2               TNAYDGKLDIFKTCRNS--       550
EptA-3               TQAYDGKLDIFKDCRTTS-       551
EptA-4               TNEYRASQDIFAPCR----       540
A. fischeri          TKAKDNALDIMAN------       542
V. cholerae El Tor   TKAYQANLDLFAKCRTSQS       550
V. vulnificus        TSEYDATLDLFKQCRAQ--       550
V. parahaemolyticus  TQAYDGQLDIFKTCRTVS-       551
                     *.      *::
```

**Fig. S3. Multiple sequence alignment of EptA-1 to -4 with EptA orthologs in *Vibrionaceae*.**

The alignment was performed using Clustal Omega (1.2.4). The conserved catalytic threonine residue (T) is in shown in red. EptA-1 to -4 sequences were chosen as one representative of each clade (amino acid sequences deduced from poolseq data). The amino acid sequences from previously characterized EptA orthologs were obtained in NCBI database and the accession numbers are: AAW87280.1 (*A. fischeri* ES114), AAF96994.1 (*V. cholerae* O1 biovar El Tor str. N16961), VVMO6_RS15980 (*V. vulnificus* MO6-24/O), and BAC62623.1 (*V. parahaemolyticus* RIMD 2210633).

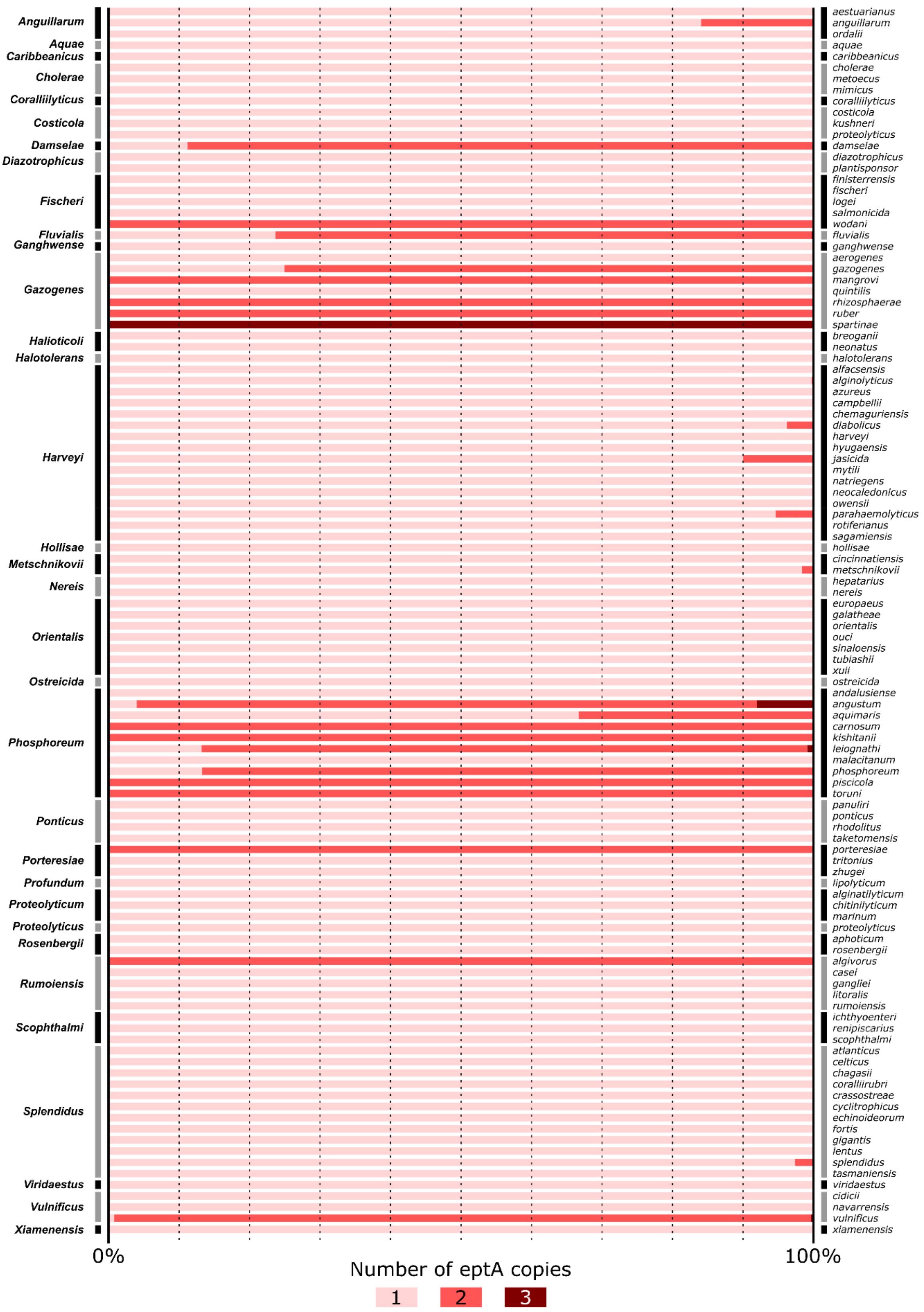


**Fig. S4. Percentage of *ept*A paralogues in species of *Vibrionaceae*.** Species of *Vibrionaceae* (right) are sorted by clade (left). The color code indicates the number of *ept*A copy per genome: pink (1 gene copy), dark pink (two gene copies), brown (3 gene copies). The scale bar indicates the percentage of genomes carrying 1, 2 or 3 *ept*A copies in each species.

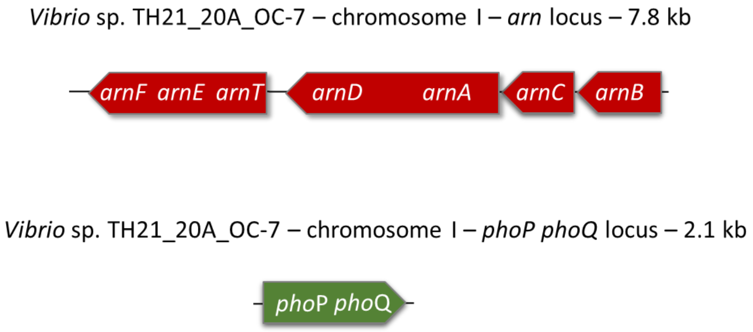


**Fig. S5. Alternative colistin resistance mechanisms identified in *Vibrio* strains TH21_20A_OC7, TH21_20_OH4, and TH21_20A_OG1**.

Each of these isolates contained both *arn*BCADTEF and phoP/Q genes within their genomes, although they were situated in distinct genomic regions.

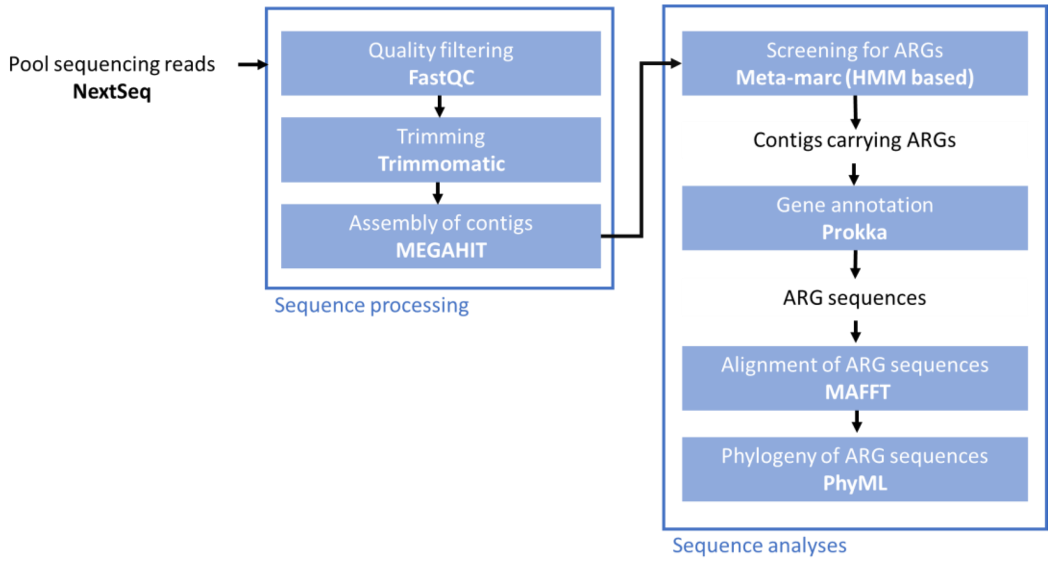


**Fig. S6. Bioinformatics analyses for identifying *ept*A and *mcr* gene sequences in Poolseq data.**

**Table S1. Metadata on *mcr/eptA* unique sequences found in Pool sequencing data**

| Sampling sites | GPS coordinates | Sampling date | Culture medium | Temp | No. CFU | Bacterial Pool ID | Total sequenced reads | Accession number | No. unique *mcr/eptA* sequences total (*mcr*; *eptA*) |
|---|---|---|---|---|---|---|---|---|---|
| **THAU** | N: 43º26.058' E: 003º39.878' | 18/10/2021 | Marine agar | 37°C | 48 | THAU-P1_5 | 7,53E+07 | SAMN37810832 | 8 (1;7) |
| | | | | 20°C | 44 | | | | |
| | | | TCBS | 37°C | 6 | THAU-P3_7 | 8,38E+07 | SAMN37810833 | 4 (0;4) |
| | | | | 20°C | 48 | | | | |
| **EBRO** | N: 40º37.106880´ E: 0º37.318320´ | 25/10/2021 | Marine agar | 37°C | 48 | EBRO-P8 | 3,59E+07 | SAMN37810834 | 6 (2;4) |
| | | | | 20°C | 47 | EBRO-P4 | 6,88E+07 | SAMN37810835 | 13 (1;12) |
| | | | TCBS | 37°C | 18 | EBRO-P6 | 1,28E+07 | SAMN37810836 | 6 (1;5) |
| | | | | 20°C | 36 | EBRO-P2 | 1,44E+07 | SAMN37810837 | **5** (0;5) |
| **SYLT** | N: 55º 1' 42.539" E: 8º 26' 1.953'' | 10/01/2022 | Marine Agar | 37°C | 32 | SYLT-P7 | 3,05E+07 | SAMN37810838 | 2 (1;1) |
| | | | | 20°C | 19 | SYLT-P1 | 6,57E+06 | SAMN37810839 | **2** (0;2) |
| | | | TCBS | 20°C | 48 | SYLT-P4 | 7,73E+07 | SAMN37810840 | 7 (4;3) |
| | | | | 37°C | 0 | na | na | na | **0** (0;0) |
| **Total** | | | | | **394** | | | | **53** (10;43) |

**Table S2. Genbank ID of nucleotide sequences of *mcr*/*ept*A genes**

| *mcr/eptA* sequence ID | Gene name | GenBank ID |
|---|---|---|
| EBRO-P2_03786 | *eptA-1* | OR578981 |
| EBRO-P6_09496 | *eptA-1* | OR578982 |
| EBRO-P6_10674 | *eptA-1* | OR578983 |
| EBRO-P2_02270 | *eptA-1* | OR578984 |
| THAU-P3-7_06040 | *eptA-1* | OR578979 |
| EBRO-P4_30301 | *eptA-1* | OR578985 |
| THAU-P1-5_20040 | *eptA-1* | OR578980 |
| EBRO-P8_20143 | *eptA-1* | OR578986 |
| EBRO-P4_33592 | *eptA-1* | OR578987 |
| EBRO-P8_12622 | *eptA-1* | OR578988 |
| EBRO-P2_00219 | *eptA-2* | OR578989 |
| EBRO-P4_18218 | *eptA-2* | OR578990 |
| EBRO-P8_23965 | *eptA-2* | OR578991 |
| EBRO-P6_03152 | *eptA-3* | OR578994 |
| EBRO-P6_08759 | *eptA-3* | OR578995 |
| THAU-P3-7_04605 | *eptA-3* | OR578992 |
| EBRO-P8_28758 | *eptA-3* | OR578996 |
| THAU-P1-5_15587 | *eptA-3* | OR578993 |
| EBRO-P2_05228 | *eptA-4* | OR579004 |
| THAU-P3-7_15942 | *eptA-4* | OR578997 |
| SYLT-P4_02614 | *eptA-4* | OR578999 |
| EBRO-P2_00375 | *eptA-4* | OR579005 |
| SYLT-P4_00252 | *eptA-4* | OR579000 |
| SYLT-P4_02050 | *eptA-4* | OR579001 |
| EBRO-P4_25590 | *eptA-4* | OR579006 |
| EBRO-P4_20292 | *eptA-4* | OR579007 |
| EBRO-P4_37349 | *eptA-4* | OR579008 |
| EBRO-P4_27855 | *eptA-4* | OR579009 |
| THAU-P1-5_47017 | *eptA-4* | OR578998 |
| EBRO-P4_30821 | *eptA-4* | OR579010 |
| SYLT-P1_02695 | *eptA-4* | OR579002 |
| SYLT-P1_02880 | *eptA-4* | OR579003 |
| SYLT-P7_16811 | other *eptA* sequence | OR579015 |
| THAU-P1-5_36515 | other *eptA* sequence | OR579011 |
| EBRO-P4_29453 | other *eptA* sequence | OR579016 |
| THAU-P1-5_40795 | other *eptA* sequence | OR579012 |
| EBRO-P4_29773 | other *eptA* sequence | OR579017 |
| EBRO-P4_25721 | other *eptA* sequence | OR579018 |
| EBRO-P6_12023 | other *eptA* sequence | OR579019 |
| EBRO-P4_14649 | other *eptA* sequence | OR579020 |
| THAU-P3-7_01996 | other *eptA* sequence | OR579013 |
| THAU-P1-5_24140 | other *eptA* sequence | OR579014 |
| SYLT-P7_12619 | other *mcr* sequence | OR579022 |

| | | |
|---|---|---|
| EBRO-P4_00710 | other *mcr* sequence | OR579026 |
| EBRO-P6_04283 | other *mcr* sequence | OR579027 |
| EBRO-P8_16485 | other *mcr* sequence | OR579028 |
| THAU-P1-5_15158 | other *mcr* sequence | OR579021 |
| SYLT-P4_10061 | other *mcr* sequence | OR579023 |
| EBRO-P8_21207 | other *mcr* sequence | OR579029 |
| SYLT-P4_06950 | other *mcr* sequence | OR579024 |
| SYLT-P4_00705 | other *mcr* sequence | OR579025 |

### Table S3. List of *mcr*/*ept*A unique sequences found in pool sequencing data

| Gene name | *mcr/eptA* sequence ID | Poolseq library | Sampling site | Genetic environment | Genus based on BlastP best hit | Culture medium | Temp | No. of mapped Reads | Normalised read counts * 10^6 |
|---|---|---|---|---|---|---|---|---|---|
| ***eptA-1*** | EBRO-P8_12622 | EBRO-P8 | EBRO | *rstA-rstB-Zipper-dgkA-eptA* | *Vibrio* | MA | 37°C | 2136 | 60,39 |
| ***eptA-1*** | THAU-P3-7_06040 | THAU-P3-7 | THAU | *rstA-rstB-Zipper-dgkA-eptA* | *Vibrio* | TCBS | 20+37°C | 7010 | 84,39 |
| ***eptA-1*** | EBRO-P2_02270 | EBRO-P2 | EBRO | *rstA-rstB-Zipper-dgkA-eptA* | *Vibrio* | TCBS | 20°C | 247 | 17,40 |
| ***eptA-1*** | EBRO-P4_33592 | EBRO-P4 | EBRO | *rstA-rstB-Zipper-dgkA-eptA* | *Vibrio* | MA | 20°C | 2440 | 35,81 |
| ***eptA-1*** | EBRO-P2_03786 | EBRO-P2 | EBRO | *rstA-rstB-Zipper-dgkA-eptA* | *Vibrio* | TCBS | 20°C | 418 | 29,44 |
| ***eptA-1*** | EBRO-P4_30301 | EBRO-P4 | EBRO | *rstA-rstB-Zipper-dgkA-eptA* | *Vibrio* | MA | 20°C | 375 | 5,50 |
| ***eptA-1*** | EBRO-P6_09496 | EBRO-P6 | EBRO | *rstA-rstB-Zipper-dgkA-eptA* | *Vibrio* | TCBS | 37°C | 356 | 28,11 |
| ***eptA-1*** | THAU-P1-5_20040 | THAU-P1-5 | THAU | *rstA-rstB-Zipper-dgkA-eptA* | *Vibrio* | MA | 20+37°C | 2669 | 35,89 |
| ***eptA-1*** | EBRO-P8_20143 | EBRO-P8 | EBRO | *rstA-rstB-Zipper-dgkA-eptA* | *Vibrio* | MA | 37°C | 1546 | 43,71 |
| ***eptA-1*** | EBRO-P6_10674 | EBRO-P6 | EBRO | *rstA-rstB-Zipper-dgkA-eptA* | *Vibrio* | TCBS | 37°C | 958 | 75,63 |
| ***eptA-2*** | EBRO-P2_00219 | EBRO-P2 | EBRO | *rstA-rstB-Zipper-dgkA-eptA* | *Vibrio* | TCBS | 20°C | 274 | 19,30 |
| ***eptA-2*** | EBRO-P4_18218 | EBRO-P4 | EBRO | *rstA-rstB-Zipper-dgkA-eptA* | *Vibrio* | MA | 20°C | 492 | 7,22 |
| ***eptA-2*** | EBRO-P8_23965 | EBRO-P8 | EBRO | *rstA-rstB-Zipper-dgkA-eptA* | *Vibrio* | MA | 37°C | 359 | 10,15 |
| ***eptA-3*** | EBRO-P8_28758 | EBRO-P8 | EBRO | *rstA-rstB-Zipper-dgkA-eptA* | *Vibrio* | MA | 37°C | 1005 | 28,41 |
| ***eptA-3*** | EBRO-P6_08759 | EBRO-P6 | EBRO | *rstA-rstB-Zipper-dgkA-eptA* | *Vibrio* | TCBS | 37°C | 333 | 26,29 |
| ***eptA-3*** | EBRO-P6_03152 | EBRO-P6 | EBRO | *rstA-rstB-Zipper-dgkA-eptA* | *Vibrio* | TCBS | 37°C | 195 | 15,40 |
| ***eptA-3*** | THAU-P1-5_15587 | THAU-P1-5 | THAU | *rstA-rstB-Zipper-dgkA-eptA* | *Vibrio* | MA | 20+37°C | 1356 | 18,23 |
| ***eptA-3*** | THAU-P3-7_04605 | THAU-P3-7 | THAU | *rstA-rstB-Zipper-dgkA-eptA* | *Vibrio* | TCBS | 20+37°C | 2751 | 33,12 |
| ***eptA-4*** | EBRO-P2_00375 | EBRO-P2 | EBRO | *eptA* | *Vibrio* | TCBS | 20°C | 256 | 18,03 |
| ***eptA-4*** | SYLT-P4_02614 | SYLT-P4 | SYLT | *eptA* | *Vibrio* | TCBS | 20°C | 156 | 2,03 |
| ***eptA-4*** | EBRO-P4_30821 | EBRO-P4 | EBRO | *eptA* | *Vibrio* | MA | 20°C | 288 | 4,23 |
| ***eptA-4*** | SYLT-P1_02695 | SYLT-P1 | SYLT | *eptA* | *Vibrio* | MA | 20°C | 1017 | 159,08 |

| | | | | | | | | | |
|---|---|---|---|---|---|---|---|---|---|
| ***eptA-4*** | SYLT-P4_00252 | SYLT-P4 | SYLT | *eptA* | *Vibrio* | TCBS | 20°C | 8823 | 115,02 |
| ***eptA-4*** | EBRO-P2_05228 | EBRO-P2 | EBRO | *eptA* | *Vibrio* | TCBS | 20°C | 753 | 53,04 |
| ***eptA-4*** | EBRO-P4_25590 | EBRO-P4 | EBRO | *eptA* | *Vibrio* | MA | 20°C | 612 | 8,98 |
| ***eptA-4*** | EBRO-P4_20292 | EBRO-P4 | EBRO | *eptA* | *Vibrio* | MA | 20°C | 946 | 13,89 |
| ***eptA-4*** | EBRO-P4_37349 | EBRO-P4 | EBRO | *eptA* | *Vibrio* | MA | 20°C | 2770 | 40,66 |
| ***eptA-4*** | SYLT-P4_02050 | SYLT-P4 | SYLT | *eptA* | *Vibrio* | TCBS | 20°C | 1745 | 22,75 |
| ***eptA-4*** | THAU-P3-7_15942 | THAU-P3-7 | THAU | *eptA* | *Vibrio* | TCBS | 20+37°C | 891 | 10,73 |
| ***eptA-4*** | EBRO-P4_27855 | EBRO-P4 | EBRO | *eptA* | *Vibrio* | MA | 20°C | 1117 | 16,40 |
| ***eptA-4*** | SYLT-P1_02880 | SYLT-P1 | SYLT | *eptA* | *Vibrio* | MA | 20°C | 1244 | 194,59 |
| ***eptA-4*** | THAU-P1-5_47017 | THAU-P1-5 | THAU | *eptA* | *Vibrio* | MA | 20+37°C | 857 | 11,52 |
| **other eptA** | SYLT-P7_16811 | SYLT-P7 | SYLT | *eptA-dgkA* | *Vibrio* | MA | 37°C | 849 | 28,11 |
| **other eptA** | EBRO-P4_25721 | EBRO-P4 | EBRO | *dgkA-eptA* | *Vibrio* | MA | 20°C | 753 | 11,05 |
| **other eptA** | THAU-P1-5_24140 | THAU-P1-5 | THAU | *dgkA-eptA* | *Vibrio* | MA | 20+37°C | 226 | 3,04 |
| **other eptA** | THAU-P3-7_01996 | THAU-P3-7 | THAU | *rstA-rstB-Zipper-eptA* | *Vibrio* | TCBS | 20+37°C | 192 | 2,31 |
| **other eptA** | THAU-P1-5_40795 | THAU-P1-5 | THAU | *eptA-arlR* | *Aliivibrio* | MA | 20+37°C | 177 | 2,38 |
| **other eptA** | EBRO-P4_14649 | EBRO-P4 | EBRO | *Zipper-eptA* | *Vibrio* | MA | 20°C | 883 | 12,96 |
| **other eptA** | THAU-P1-5_36515 | THAU-P1-5 | THAU | *eptA-PGT1-arlR-sasA* | *Photobacterium* | MA | 20+37°C | 230 | 3,09 |
| **other eptA** | THAU-P1-5_32984 | THAU-P1-5 | THAU | *eptA* | *Photobacterium* | MA | 20+37°C | 195 | 2,62 |
| **other eptA** | EBRO-P4_29453 | EBRO-P4 | EBRO | *eptA* | *Aliivibrio* | MA | 20°C | 953 | 13,99 |
| ***other eptA*** | SYLT-P7_12619 | SYLT-P7 | SYLT | *dgkA-hpap2-eptA* | *Halomonas* | MA | 37°C | 954 | 31,58 |
| **other eptA** | EBRO-P6_12023 | EBRO-P6 | EBRO | *dgkA-eptA* | *Vibrio* | TCBS | 37°C | 225 | 17,76 |
| **other eptA** | EBRO-P4_29773 | EBRO-P4 | EBRO | *dgkA-eptA-PGT1* | *Vibrio* | MA | 20°C | 373 | 5,47 |
| ***other mcr*** | EBRO-P4_00710 | EBRO-P4 | EBRO | *mcr* | *Vibrio* | MA | 20°C | 580 | 8,51 |
| ***other mcr*** | SYLT-P4_10061 | SYLT-P4 | SYLT | *mcr-dgkA* | *Photobacterium* | TCBS | 20°C | 517 | 6,74 |
| ***other mcr*** | SYLT-P4_06950 | SYLT-P4 | SYLT | *mcr* | *Shewanella* | TCBS | 20°C | 754 | 9,83 |
| ***other mcr*** | SYLT-P4_13141 | SYLT-P4 | SYLT | *mcr* | *Shewanella* | TCBS | 20°C | 112 | 1,46 |
| ***other mcr*** | EBRO-P6_04283 | EBRO-P6 | EBRO | *mcr-dgkA* | *Vibrio* | TCBS | 37°C | 211 | 16,66 |
| ***other mcr*** | THAU-P1-5_15158 | THAU-P1-5 | THAU | *mcr-dgkA* | *Vibrio* | MA | 20+37°C | 292 | 3,93 |
| ***other mcr*** | EBRO-P8_21207 | EBRO-P8 | EBRO | *mcr-dgkA* | *Pseudoalteromonas* | MA | 37°C | 154 | 4,35 |
| ***other mcr*** | SYLT-P4_00705 | SYLT-P4 | SYLT | *mcr* | *Shewanella* | TCBS | 20°C | 386 | 5,03 |
| ***other mcr*** | EBRO-P8_16485 | EBRO-P8 | EBRO | *mcr-dgkA* | *Vibrio* | MA | 37°C | 721 | 20,38 |

### Table S4. Amino acid sequence identity between EptA variants

| | **VF_A0210** | EBRO-P4_29453\|OR579016\|eptA-like | THAU-P1-5_40795\|OR579012\|eptA-like | SYLT-P7_16811\|OR579015\|eptA-like | THAU-P1-5_36515\|OR579011\|eptA-like | SYLT-P1_02880\|OR579003\|eptA-4 | EBRO-P4_25590\|OR579006\|eptA-4 | SYLT-P4_02050\|OR579001\|eptA-4 | EBRO-P2_00375\|OR579005\|eptA-4 | SYLT-P1_02695\|OR579002\|eptA-4 | EBRO-P4_30821\|OR579010\|eptA-4 | SYLT-P4_00252\|OR579000\|eptA-4 | EBRO-P4_20292\|OR579007\|eptA-4 | EBRO-P2_05228\|OR579004\|eptA-4 | EBRO-P4_37349\|OR579008\|eptA-4 | SYLT-P4_02614\|OR578999\|eptA-4 | THAU-P3-7_15942\|OR578997\|eptA-4 | EBRO-P4_27855\|OR579009\|eptA-4 | THAU-P1-5_47017\|OR578998\|eptA-4 | EBRO-P2_00219\|OR578989\|eptA-2 | EBRO-P4_18218\|OR578990\|eptA-2 | EBRO-P8_23965\|OR578991\|eptA-2 | EBRO-P2_03786\|OR578981\|eptA-1 | THAU-P1-5_20040\|OR578980\|eptA-1 | EBRO-P4_30301\|OR578985\|eptA-1 | EBRO-P6_09496\|OR578982\|eptA-1 | EBRO-P6_10674\|OR578983\|eptA-1 | EBRO-P8_20143\|OR578986\|eptA-1 | EBRO-P2_02270\|OR578984\|eptA-1 | EBRO-P4_33592\|OR578987\|eptA-1 | EBRO-P8_12622\|OR578988\|eptA-1 | THAU-P3-7_06040\|OR578979\|eptA-1 | EBRO-P6_08759\|OR578995\|eptA-3 | THAU-P1-5_15587\|OR578993\|eptA-3 | THAU-P3-7_04605\|OR578992\|eptA-3 | EBRO-P6_03152\|OR578994\|eptA-3 | EBRO-P8_28758\|OR578996\|eptA-3 | THAU-P1-5_24140\|OR579014\|eptA-like | THAU-P3-7_01996\|OR579013\|eptA-like | **VP_RS21300** | **VVMO6_RS15980** | **VCA1102** | EBRO-P4_14649\|OR579020\|eptA-like | EBRO-P6_12023\|OR579019\|eptA-like | EBRO-P4_25721\|OR579018\|eptA-like | EBRO-P4_29773\|OR579017\|eptA-like |
|---|---|---|---|---|---|---|---|---|---|---|---|---|---|---|---|---|---|---|---|---|---|---|---|---|---|---|---|---|---|---|---|---|---|---|---|---|---|---|---|---|---|---|---|---|---|---|
| **VF_A0210** | 100 | 99 | 99 | 51 | 51 | 44 | 44 | 44 | 43 | 43 | 43 | 43 | 43 | 43 | 43 | 43 | 43 | 43 | 43 | 44 | 44 | 44 | 44 | 44 | 43 | 43 | 44 | 44 | 44 | 44 | 44 | 44 | 43 | 42 | 42 | 42 | 42 | 41 | 44 | 43 | 43 | 43 | 40 | 43 | 44 | 42 |
| EBRO-P4_29453\|OR579016\|eptA-like | 99 | 100 | 99 | 51 | 51 | 44 | 44 | 44 | 43 | 43 | 43 | 43 | 43 | 43 | 43 | 43 | 43 | 43 | 43 | 45 | 45 | 45 | 44 | 44 | 44 | 44 | 44 | 44 | 44 | 44 | 44 | 44 | 43 | 43 | 43 | 43 | 42 | 41 | 44 | 43 | 43 | 43 | 41 | 44 | 44 | 42 |
| THAU-P1-5_40795\|OR579012\|eptA-like | 99 | 99 | 100 | 52 | 51 | 44 | 44 | 44 | 43 | 43 | 43 | 43 | 43 | 43 | 43 | 43 | 43 | 43 | 43 | 45 | 45 | 45 | 44 | 44 | 44 | 44 | 44 | 44 | 44 | 44 | 44 | 44 | 43 | 43 | 43 | 43 | 42 | 41 | 44 | 43 | 43 | 43 | 41 | 44 | 44 | 42 |
| SYLT-P7_16811\|OR579015\|eptA-like | 51 | 51 | 52 | 100 | 54 | 43 | 43 | 43 | 43 | 43 | 43 | 43 | 43 | 43 | 44 | 42 | 43 | 43 | 42 | 45 | 45 | 45 | 44 | 44 | 44 | 44 | 44 | 44 | 45 | 45 | 44 | 44 | 42 | 42 | 42 | 42 | 42 | 41 | 44 | 43 | 44 | 41 | 41 | 41 | 42 | 42 |
| THAU-P1-5_36515\|OR579011\|eptA-like | 51 | 51 | 51 | 54 | 100 | 43 | 43 | 43 | 43 | 43 | 43 | 43 | 41 | 41 | 41 | 41 | 42 | 42 | 42 | 45 | 45 | 45 | 45 | 44 | 44 | 44 | 44 | 44 | 44 | 44 | 44 | 44 | 42 | 42 | 42 | 43 | 43 | 44 | 44 | 43 | 44 | 42 | 42 | 43 | 43 | 41 |
| SYLT-P1_02880\|OR579003\|eptA-4 | 44 | 44 | 44 | 43 | 43 | 100 | 100 | 100 | 94 | 94 | 94 | 94 | 86 | 87 | 86 | 88 | 90 | 89 | 89 | 53 | 53 | 53 | 53 | 53 | 53 | 53 | 53 | 53 | 53 | 53 | 53 | 52 | 53 | 53 | 53 | 53 | 53 | 52 | 54 | 54 | 55 | 54 | 50 | 51 | 50 | 50 |
| EBRO-P4_25590\|OR579006\|eptA-4 | 44 | 44 | 44 | 43 | 43 | 100 | 100 | 100 | 94 | 94 | 94 | 94 | 86 | 86 | 86 | 88 | 89 | 89 | 88 | 53 | 53 | 53 | 53 | 53 | 53 | 53 | 53 | 53 | 53 | 53 | 52 | 52 | 52 | 52 | 52 | 53 | 52 | 52 | 54 | 54 | 55 | 54 | 50 | 51 | 50 | 50 |
| SYLT-P4_02050\|OR579001\|eptA-4 | 44 | 44 | 44 | 43 | 43 | 100 | 100 | 100 | 94 | 94 | 94 | 94 | 86 | 86 | 86 | 88 | 89 | 89 | 88 | 53 | 53 | 53 | 53 | 53 | 53 | 53 | 53 | 53 | 53 | 53 | 52 | 52 | 52 | 52 | 52 | 53 | 52 | 52 | 54 | 54 | 55 | 54 | 50 | 51 | 50 | 50 |
| EBRO-P2_00375\|OR579005\|eptA-4 | 43 | 43 | 43 | 43 | 43 | 94 | 94 | 94 | 100 | 97 | 97 | 97 | 87 | 88 | 87 | 88 | 90 | 90 | 90 | 53 | 53 | 53 | 54 | 54 | 54 | 54 | 54 | 54 | 54 | 54 | 53 | 53 | 53 | 53 | 53 | 54 | 54 | 54 | 55 | 54 | 56 | 55 | 50 | 51 | 51 | 50 |
| SYLT-P1_02695\|OR579002\|eptA-4 | 43 | 43 | 43 | 43 | 43 | 94 | 94 | 94 | 97 | 100 | 99 | 99 | 87 | 88 | 88 | 88 | 90 | 91 | 91 | 54 | 54 | 54 | 54 | 54 | 54 | 54 | 54 | 54 | 54 | 54 | 53 | 53 | 54 | 54 | 54 | 54 | 54 | 54 | 55 | 55 | 56 | 55 | 51 | 51 | 51 | 51 |
| EBRO-P4_30821\|OR579010\|eptA-4 | 43 | 43 | 43 | 43 | 43 | 94 | 94 | 94 | 97 | 99 | 100 | 100 | 87 | 87 | 87 | 88 | 90 | 91 | 91 | 54 | 54 | 54 | 54 | 54 | 54 | 54 | 54 | 54 | 54 | 54 | 54 | 53 | 54 | 54 | 54 | 54 | 54 | 54 | 55 | 55 | 57 | 55 | 51 | 52 | 51 | 51 |
| SYLT-P4_00252\|OR579000\|eptA-4 | 43 | 43 | 43 | 43 | 43 | 94 | 94 | 94 | 97 | 99 | 100 | 100 | 87 | 87 | 87 | 88 | 90 | 91 | 91 | 54 | 54 | 54 | 54 | 54 | 54 | 54 | 54 | 54 | 54 | 54 | 54 | 53 | 54 | 54 | 54 | 54 | 54 | 54 | 55 | 55 | 57 | 55 | 51 | 52 | 51 | 51 |
| EBRO-P4_20292\|OR579007\|eptA-4 | 43 | 43 | 43 | 43 | 41 | 86 | 86 | 86 | 87 | 87 | 87 | 87 | 100 | 94 | 94 | 90 | 90 | 91 | 91 | 52 | 52 | 52 | 52 | 52 | 52 | 52 | 53 | 53 | 52 | 52 | 51 | 51 | 52 | 52 | 52 | 52 | 52 | 52 | 53 | 53 | 55 | 55 | 49 | 50 | 49 | 50 |
| EBRO-P2_05228\|OR579004\|eptA-4 | 43 | 43 | 43 | 43 | 41 | 87 | 86 | 86 | 88 | 88 | 87 | 87 | 94 | 100 | 99 | 93 | 91 | 92 | 91 | 53 | 53 | 53 | 53 | 53 | 53 | 53 | 54 | 54 | 53 | 53 | 52 | 52 | 52 | 52 | 52 | 52 | 52 | 53 | 53 | 54 | 56 | 55 | 49 | 51 | 49 | 50 |
| EBRO-P4_37349\|OR579008\|eptA-4 | 43 | 43 | 43 | 44 | 41 | 86 | 86 | 86 | 87 | 88 | 87 | 87 | 94 | 99 | 100 | 93 | 91 | 91 | 91 | 52 | 52 | 52 | 53 | 53 | 53 | 53 | 53 | 53 | 53 | 53 | 52 | 52 | 52 | 52 | 52 | 52 | 52 | 52 | 53 | 53 | 56 | 55 | 49 | 51 | 49 | 50 |
| SYLT-P4_02614\|OR578999\|eptA-4 | 43 | 43 | 43 | 42 | 41 | 88 | 88 | 88 | 88 | 88 | 88 | 88 | 90 | 93 | 93 | 100 | 94 | 93 | 92 | 52 | 52 | 52 | 52 | 52 | 52 | 52 | 53 | 53 | 53 | 53 | 52 | 52 | 54 | 54 | 54 | 54 | 53 | 52 | 53 | 55 | 56 | 55 | 49 | 52 | 49 | 50 |
| THAU-P3-7_15942\|OR578997\|eptA-4 | 43 | 43 | 43 | 43 | 42 | 90 | 89 | 89 | 90 | 90 | 90 | 90 | 90 | 91 | 91 | 94 | 100 | 94 | 93 | 53 | 53 | 53 | 53 | 53 | 53 | 53 | 54 | 54 | 54 | 54 | 54 | 53 | 53 | 53 | 53 | 53 | 52 | 52 | 54 | 54 | 56 | 54 | 49 | 51 | 49 | 49 |
| EBRO-P4_27855\|OR579009\|eptA-4 | 43 | 43 | 43 | 43 | 42 | 89 | 89 | 89 | 90 | 91 | 91 | 91 | 91 | 92 | 91 | 93 | 94 | 100 | 96 | 52 | 52 | 52 | 53 | 53 | 53 | 53 | 53 | 53 | 52 | 52 | 53 | 52 | 53 | 53 | 53 | 53 | 53 | 53 | 54 | 54 | 56 | 55 | 50 | 52 | 50 | 50 |
| THAU-P1-5_47017\|OR578998\|eptA-4 | 43 | 43 | 43 | 42 | 42 | 89 | 88 | 88 | 90 | 91 | 91 | 91 | 91 | 91 | 91 | 92 | 93 | 96 | 100 | 52 | 52 | 52 | 52 | 52 | 52 | 52 | 53 | 53 | 52 | 52 | 52 | 52 | 54 | 54 | 54 | 53 | 53 | 52 | 54 | 54 | 55 | 54 | 49 | 52 | 50 | 50 |
| EBRO-P2_00219\|OR578989\|eptA-2 | 44 | 45 | 45 | 45 | 45 | 53 | 53 | 53 | 53 | 54 | 54 | 54 | 52 | 53 | 52 | 52 | 53 | 52 | 52 | 100 | 100 | 99 | 87 | 88 | 87 | 87 | 88 | 88 | 87 | 87 | 87 | 87 | 79 | 79 | 78 | 79 | 79 | 76 | 82 | 84 | 73 | 59 | 62 | 65 | 61 | 62 |
| EBRO-P4_18218\|OR578990\|eptA-2 | 44 | 45 | 45 | 45 | 45 | 53 | 53 | 53 | 53 | 54 | 54 | 54 | 52 | 53 | 52 | 52 | 53 | 52 | 52 | 100 | 100 | 100 | 87 | 87 | 87 | 87 | 87 | 87 | 86 | 87 | 87 | 86 | 78 | 78 | 78 | 79 | 78 | 76 | 82 | 83 | 73 | 59 | 62 | 65 | 61 | 62 |
| EBRO-P8_23965\|OR578991\|eptA-2 | 44 | 45 | 45 | 45 | 45 | 53 | 53 | 53 | 53 | 54 | 54 | 54 | 52 | 53 | 52 | 52 | 53 | 52 | 52 | 99 | 100 | 100 | 87 | 87 | 87 | 87 | 87 | 87 | 86 | 87 | 87 | 87 | 78 | 78 | 78 | 79 | 78 | 76 | 82 | 83 | 73 | 59 | 62 | 65 | 61 | 62 |
| EBRO-P2_03786\|OR578981\|eptA-1 | 44 | 44 | 44 | 44 | 45 | 53 | 53 | 53 | 54 | 54 | 54 | 54 | 52 | 53 | 53 | 52 | 53 | 53 | 52 | 87 | 87 | 87 | 100 | 100 | 99 | 99 | 92 | 92 | 91 | 91 | 92 | 92 | 77 | 77 | 77 | 77 | 77 | 76 | 81 | 80 | 70 | 58 | 61 | 64 | 62 | 62 |
| THAU-P1-5_20040\|OR578980\|eptA-1 | 44 | 44 | 44 | 44 | 44 | 53 | 53 | 53 | 54 | 54 | 54 | 54 | 52 | 53 | 53 | 52 | 53 | 53 | 52 | 88 | 87 | 87 | 100 | 100 | 100 | 100 | 92 | 92 | 91 | 91 | 92 | 92 | 77 | 77 | 77 | 77 | 77 | 76 | 81 | 80 | 71 | 58 | 62 | 64 | 62 | 62 |
| EBRO-P4_30301\|OR578985\|eptA-1 | 43 | 44 | 44 | 44 | 44 | 53 | 53 | 53 | 54 | 54 | 54 | 54 | 52 | 53 | 53 | 52 | 53 | 53 | 52 | 87 | 87 | 87 | 99 | 100 | 100 | 100 | 92 | 92 | 91 | 91 | 92 | 92 | 77 | 77 | 77 | 77 | 76 | 76 | 81 | 80 | 70 | 58 | 61 | 63 | 62 | 61 |
| EBRO-P6_09496\|OR578982\|eptA-1 | 43 | 44 | 44 | 44 | 44 | 53 | 53 | 53 | 54 | 54 | 54 | 54 | 52 | 53 | 53 | 52 | 53 | 53 | 52 | 87 | 87 | 87 | 99 | 100 | 100 | 100 | 92 | 92 | 91 | 91 | 92 | 92 | 77 | 77 | 77 | 77 | 76 | 76 | 81 | 80 | 70 | 58 | 61 | 63 | 62 | 61 |
| EBRO-P6_10674\|OR578983\|eptA-1 | 44 | 44 | 44 | 44 | 44 | 53 | 53 | 53 | 54 | 54 | 54 | 54 | 53 | 54 | 53 | 53 | 54 | 53 | 53 | 88 | 87 | 87 | 92 | 92 | 92 | 92 | 100 | 99 | 93 | 93 | 94 | 94 | 78 | 78 | 78 | 79 | 79 | 77 | 81 | 81 | 71 | 58 | 61 | 63 | 63 | 61 |
| EBRO-P8_20143\|OR578986\|eptA-1 | 44 | 44 | 44 | 44 | 44 | 53 | 53 | 53 | 54 | 54 | 54 | 54 | 53 | 54 | 53 | 53 | 54 | 53 | 53 | 88 | 87 | 87 | 92 | 92 | 92 | 92 | 99 | 100 | 93 | 93 | 95 | 94 | 78 | 78 | 78 | 79 | 78 | 77 | 81 | 81 | 71 | 58 | 61 | 63 | 62 | 61 |
| EBRO-P2_02270\|OR578984\|eptA-1 | 44 | 44 | 44 | 45 | 44 | 53 | 53 | 53 | 54 | 54 | 54 | 54 | 52 | 53 | 53 | 53 | 54 | 52 | 52 | 87 | 86 | 86 | 91 | 91 | 91 | 91 | 93 | 93 | 100 | 100 | 95 | 95 | 77 | 77 | 77 | 77 | 77 | 77 | 81 | 81 | 69 | 58 | 61 | 63 | 61 | 60 |
| EBRO-P4_33592\|OR578987\|eptA-1 | 44 | 44 | 44 | 45 | 44 | 53 | 53 | 53 | 54 | 54 | 54 | 54 | 52 | 53 | 53 | 53 | 54 | 52 | 52 | 87 | 87 | 87 | 91 | 91 | 91 | 91 | 93 | 93 | 100 | 100 | 95 | 95 | 77 | 77 | 77 | 77 | 77 | 77 | 81 | 81 | 69 | 58 | 61 | 63 | 61 | 60 |
| EBRO-P8_12622\|OR578988\|eptA-1 | 44 | 44 | 44 | 44 | 44 | 53 | 52 | 52 | 53 | 53 | 54 | 54 | 51 | 52 | 52 | 52 | 54 | 53 | 52 | 87 | 87 | 87 | 92 | 92 | 92 | 92 | 94 | 95 | 95 | 95 | 100 | 99 | 77 | 77 | 77 | 77 | 77 | 78 | 81 | 80 | 70 | 58 | 61 | 63 | 61 | 61 |
| THAU-P3-7_06040\|OR578979\|eptA-1 | 44 | 44 | 44 | 44 | 44 | 52 | 52 | 52 | 53 | 53 | 53 | 53 | 51 | 52 | 52 | 52 | 53 | 52 | 52 | 87 | 86 | 87 | 92 | 92 | 92 | 92 | 94 | 94 | 95 | 95 | 99 | 100 | 77 | 77 | 77 | 78 | 77 | 77 | 80 | 80 | 70 | 57 | 61 | 63 | 61 | 60 |
| EBRO-P6_08759\|OR578995\|eptA-3 | 43 | 43 | 43 | 42 | 42 | 53 | 52 | 52 | 53 | 54 | 54 | 54 | 52 | 52 | 52 | 54 | 53 | 53 | 54 | 79 | 78 | 78 | 77 | 77 | 77 | 77 | 78 | 78 | 77 | 77 | 77 | 77 | 100 | 99 | 99 | 92 | 91 | 77 | 77 | 83 | 70 | 57 | 60 | 63 | 59 | 63 |
| THAU-P1-5_15587\|OR578993\|eptA-3 | 42 | 43 | 43 | 42 | 42 | 53 | 52 | 52 | 53 | 54 | 54 | 54 | 52 | 52 | 52 | 54 | 53 | 53 | 54 | 79 | 78 | 78 | 77 | 77 | 77 | 77 | 78 | 78 | 77 | 77 | 77 | 77 | 99 | 100 | 100 | 92 | 91 | 77 | 77 | 83 | 70 | 57 | 60 | 63 | 59 | 63 |
| THAU-P3-7_04605\|OR578992\|eptA-3 | 42 | 43 | 43 | 42 | 42 | 53 | 52 | 52 | 53 | 54 | 54 | 54 | 52 | 52 | 52 | 54 | 53 | 53 | 54 | 78 | 78 | 78 | 77 | 77 | 77 | 77 | 78 | 78 | 77 | 77 | 77 | 77 | 99 | 100 | 100 | 92 | 91 | 77 | 77 | 83 | 70 | 57 | 60 | 62 | 59 | 63 |
| EBRO-P6_03152\|OR578994\|eptA-3 | 42 | 43 | 43 | 42 | 43 | 53 | 53 | 53 | 54 | 54 | 54 | 54 | 52 | 52 | 52 | 54 | 53 | 53 | 53 | 79 | 79 | 79 | 77 | 77 | 77 | 77 | 79 | 79 | 77 | 77 | 77 | 78 | 92 | 92 | 92 | 100 | 99 | 78 | 78 | 84 | 70 | 58 | 61 | 63 | 58 | 62 |
| EBRO-P8_28758\|OR578996\|eptA-3 | 42 | 42 | 42 | 42 | 43 | 53 | 52 | 52 | 54 | 54 | 54 | 54 | 52 | 52 | 52 | 53 | 52 | 53 | 53 | 79 | 78 | 78 | 77 | 77 | 76 | 76 | 79 | 78 | 77 | 77 | 77 | 77 | 91 | 91 | 91 | 99 | 100 | 78 | 78 | 83 | 70 | 58 | 60 | 63 | 58 | 62 |
| THAU-P1-5_24140\|OR579014\|eptA-like | 41 | 41 | 41 | 41 | 44 | 52 | 52 | 52 | 54 | 54 | 54 | 54 | 52 | 53 | 52 | 52 | 52 | 53 | 52 | 76 | 76 | 76 | 76 | 76 | 76 | 76 | 77 | 77 | 77 | 77 | 78 | 77 | 77 | 77 | 77 | 78 | 78 | 100 | 76 | 79 | 69 | 57 | 60 | 61 | 60 | 61 |
| THAU-P3-7_01996\|OR579013\|eptA-like | 44 | 44 | 44 | 44 | 44 | 54 | 54 | 54 | 55 | 55 | 55 | 55 | 53 | 53 | 53 | 53 | 54 | 54 | 54 | 82 | 82 | 82 | 81 | 81 | 81 | 81 | 81 | 81 | 81 | 81 | 81 | 80 | 77 | 77 | 77 | 78 | 78 | 76 | 100 | 80 | 69 | 57 | 61 | 64 | 61 | 61 |
| **VP_RS21300** | 43 | 43 | 43 | 43 | 43 | 54 | 54 | 54 | 54 | 55 | 55 | 55 | 53 | 54 | 53 | 55 | 54 | 54 | 54 | 84 | 83 | 83 | 80 | 80 | 80 | 80 | 81 | 81 | 81 | 81 | 80 | 80 | 83 | 83 | 83 | 84 | 83 | 79 | 80 | 100 | 70 | 58 | 61 | 64 | 61 | 62 |
| **VVMO6_RS15980** | 43 | 43 | 43 | 44 | 44 | 55 | 55 | 55 | 56 | 56 | 57 | 57 | 55 | 56 | 56 | 56 | 56 | 56 | 55 | 73 | 73 | 73 | 70 | 71 | 70 | 70 | 71 | 71 | 69 | 69 | 70 | 70 | 70 | 70 | 70 | 70 | 70 | 69 | 69 | 70 | 100 | 60 | 62 | 63 | 62 | 66 |
| **VCA1102** | 43 | 43 | 43 | 41 | 42 | 54 | 54 | 54 | 55 | 55 | 55 | 55 | 55 | 55 | 55 | 55 | 54 | 55 | 54 | 59 | 59 | 59 | 58 | 58 | 58 | 58 | 58 | 58 | 58 | 58 | 58 | 57 | 57 | 57 | 57 | 58 | 58 | 57 | 57 | 58 | 60 | 100 | 54 | 53 | 55 | 57 |
| EBRO-P4_14649\|OR579020\|eptA-like | 40 | 41 | 41 | 41 | 42 | 50 | 50 | 50 | 50 | 51 | 51 | 51 | 49 | 49 | 49 | 49 | 49 | 50 | 49 | 62 | 62 | 62 | 61 | 62 | 61 | 61 | 61 | 61 | 61 | 61 | 61 | 61 | 60 | 60 | 60 | 61 | 60 | 60 | 61 | 61 | 62 | 54 | 100 | 64 | 57 | 56 |
| EBRO-P6_12023\|OR579019\|eptA-like | 43 | 44 | 44 | 41 | 43 | 51 | 51 | 51 | 51 | 51 | 52 | 52 | 50 | 51 | 51 | 52 | 51 | 52 | 52 | 65 | 65 | 65 | 64 | 64 | 63 | 63 | 63 | 63 | 63 | 63 | 63 | 63 | 63 | 63 | 62 | 63 | 63 | 61 | 64 | 64 | 63 | 53 | 64 | 100 | 55 | 57 |
| EBRO-P4_25721\|OR579018\|eptA-like | 44 | 44 | 44 | 42 | 43 | 50 | 50 | 50 | 51 | 51 | 51 | 51 | 49 | 49 | 49 | 49 | 49 | 50 | 50 | 61 | 61 | 61 | 62 | 62 | 62 | 62 | 63 | 62 | 61 | 61 | 61 | 61 | 59 | 59 | 59 | 58 | 58 | 60 | 61 | 61 | 62 | 55 | 57 | 55 | 100 | 58 |
| EBRO-P4_29773\|OR579017\|eptA-like | 42 | 42 | 42 | 42 | 41 | 50 | 50 | 50 | 50 | 51 | 51 | 51 | 50 | 50 | 50 | 50 | 49 | 50 | 50 | 62 | 62 | 62 | 62 | 62 | 61 | 61 | 61 | 61 | 60 | 60 | 61 | 60 | 63 | 63 | 63 | 62 | 62 | 61 | 61 | 62 | 66 | 57 | 56 | 57 | 58 | 100 |

| **% identitty** | 40 | 50 | 60 | 70 | 80 | 90 | ## |
|---|---|---|---|---|---|---|---|

**Table S5. Strains and Plasmids**

| Bacterial strains | Superclade | Isolation Source | Temp | Description | Reference | Accession Number | Conting number | size (Mb) | CDSs | Coverage |
|---|---|---|---|---|---|---|---|---|---|---|
| *Vibrio harveyi* Th15_Z_F11 | Harveyi | Water column, Thau lagoon, France | 20ºC | eptA-1; [Col-R] | (Oyanedel et al., 2023) | ERS9919775 | 53 | 5,7 | 5395 | |
| *Vibrio harveyi* Th15_F5_F11 | Harveyi | Water column, Thau lagoon, France | 20ºC | eptA-1; [Col-R] | (Oyanedel et al., 2023) | ERS9919777 | 106 | 5,9 | 5524 | |
| *Vibrio jasicida* Th15_F5_H11 | Harveyi | Water column, Thau lagoon, France | 20ºC | eptA-1; [Col-R] | (Oyanedel et al., 2023) | ERS9919787 | 52 | 6,1 | 5712 | |
| *Vibrio owensii* Th15_Z_G08 | Harveyi | Water column, Thau lagoon, France | 20ºC | eptA-1; [Col-R] | (Oyanedel et al., 2023) | ERS9919782 | 98 | 6,0 | 5566 | |
| *Vibrio rotiferianus* Th15_O_H08 | Harveyi | Oyster, Thau lagoon, France | 20ºC | eptA-2; [Col-R] | (Oyanedel et al., 2023) | ERS9919779 | 81 | 5,2 | 4835 | |
| *Vibrio rotiferianus* Th15_O_G05 | Harveyi | Oyster, Thau lagoon, France | 20ºC | eptA-2; [Col-R] | (Oyanedel et al., 2023) | ERS9919778 | 85 | 5,3 | 4956 | |
| *Vibrio splendidus* 7T7_2 | Harveyi | Oyster, Bay of Brest, France | 20ºC | eptA-4; [Col-S] | (Bruto et al., 2016) | | 297 | 5,9 | 5594 | |
| *Vibrio crassostreae* J2-4 | Splendidus | Oyster, Bay of Brest, France | 20ºC | eptA-4; [Col-S] | (Lemire et al., 2015) | PRJEB5888 | 84 | 5,7 | 5285 | |
| *Vibrio crassostreae* J2-14 | Splendidus | Oyster, Bay of Brest, France | 20ºC | eptA-4; [Col-S] | (Lemire et al., 2015) | PRJEB5892 | 65 | 5,7 | 5244 | |
| *Vibrio crassostreae* J5-15 | Splendidus | Oyster, Bay of Brest, France | 20ºC | eptA-4; [Col-S] | (Lemire et al., 2015) | PRJEB5879 | 92 | 5,7 | 5345 | |
| *Vibrio sp.* TH21_20_OG1 | unknown | Oyster, Thau lagoon, France | 20ºC | eptA-, mcr-; [Col-R] | this study | ERR12116512 | 56 | 5,4 | 4945 | 78X |
| *Vibrio sp.* TH21_20_OH4 | unknown | Oyster, Thau lagoon, France | 20ºC | eptA-, mcr-; [Col-R] | this study | ERR12116510 | 76 | 5,4 | 4903 | 34X |
| *Vibrio sp.* TH21_20A_OC7 | unknown | Oyster, Thau lagoon, France | 20ºC | eptA-, mcr-; [Col-R] | this study | ERR12116511 | 76 | 5,4 | 4790 | 68X |

| | | | | | | | | | | |
|---|---|---|---|---|---|---|---|---|---|---|
| *Vibrio sp.* TH21_20A_OB7 | Harveyi | Oyster, Thau lagoon, France | 20ºC | eptA-1; [Col-R] | this study | ERR12116517 | 177 | 6 | 5667 | 8X |
| *Vibrio jasicida* TH21_20A_OE8 | Harveyi | Oyster, Thau lagoon, France | 20ºC | eptA-1; [Col-R] | this study | ERR12116518 | 215 | 6,3 | 5824 | 33X |
| *Vibrio owensii* TH21_37_OE7 | Harveyi | Oyster, Thau lagoon, France | 37ºC | eptA-1; [Col-R] | this study | ERR12116513 | 71 | 5,9 | 5438 | 47X |
| *Vibrio alginolyticus* TH21_37_OE9 | Harveyi | Oyster, Thau lagoon, France | 37ºC | eptA-3; [Col-R] | this study | ERR12116514 | 68 | 5,3 | 4893 | 54X |
| *Vibrio alginolyticus* TH21_37A_OE10 | Harveyi | Oyster, Thau lagoon, France | 37ºC | eptA-3; [Col-R] | this study | ERR12116515 | 75 | 5,1 | 4690 | 53X |
| *Vibrio alginolyticus* TH21_37A_OE12 | Harveyi | Oyster, Thau lagoon, France | 37ºC | eptA-3; [Col-R] | this study | ERR12116516 | 88 | 5,3 | 4877 | 25X |
| *Vibrio harveyi* TH15_F5_F11 Δ*rstA* | Harveyi | na | 20ºC | *DrstA* | this study | | | | | |
| *Vibrio harveyi* TH15_F5_F11 Δ*rstA* pMRB-P$_{LAC}$-rstA | Harveyi | na | 20ºC | Δ*rstA, rstA* under plac promoter in pMRB plasmid | this study | | | | | |
| *Vibrio harvey*i TH15_F5_F11 Δ*rstA* pMRB-P$_{LAC}$-gfp | Harveyi | na | 20ºC | Δ*rstA, gfp* under plac promoter in pMRB plasmid | this study | | | | | |
| *E. coli* P3813 | | na | 37ºC | *lacIQ, thi1, supE44, endA1, recA1, hsdR17, gyrA462, zei298::Tn10,* D*thyA::(erm-pir116)* [Tc$^{R}$ Erm $^{R}$] | (Le Roux et al., 2007) | | | | | |
| *E. coli* b3914 | | na | 37ºC | (F $^{-}$) RP4-2-Tc::Mu DdapA ::*(erm-pir116), gyrA462,* | (Le Roux et al., 2007) | | | | | |

| | | | | | |
|---|---|---|---|---|---|
| | | | | *zei298::Tn10* [$Km^R$ $Em^R$ $Tc^R$] | |
| *E. coli* TOP10 | | na | 37ºC | TOP10 Chemically Competent E. coli | Invitrogen |
| *Vibrio parahaemolyticus* IFVp22 | Harveyi | Mussel, English channel, France | 37ºC | eptA; [Col-R] | (Sorée M. et al., 2024) |
| *Vibrio vulnificus* CECT4999 | Vulnificus | Eel, Spain | 37ºC | eptA; [Col-R] | (Murciano C. et al., 2017) |

| **Plasmids** | **Description** | **Reference** |
|---|---|---|
| pSW7848T | *oriV*$_{R6K}$ ; *oriT*$_{RP4}$; *araC*-P$_{BAD}$*ccdB;* [CmR] | (Val et al., 2012) |
| pSW7848 Δ*rstA* | pSW7848T :: ΔrstA | this study |
| pMRB-P$_{LAC}$-*gfp* | *oriVR6Kg ; oriTRP4; oriVpB1067;* P$_{LAC}$-*gfp* [CmR] | (Le Roux et al., 2011) |
| pMRB-P$_{LAC}$-rstA | *oriVR6Kg ; oriTRP4; oriVpB1067*P$_{LAC}$-rstA [CmR] | this study |
| pBAD-TOPO | pBAD TOPO® TA Expression Kit (Catalog nos. K4300-01, K4300-40) | Invitrogen |
| pBAD-TOPO-*dgkA-eptA-1* | *dgkA-eptA-1* under pBAD promoter in pBAD TOPO® TA | this study |
| pBAD-TOPO-*eptA-1* | *eptA-1* under pBAD promoter in pBAD TOPO® TA | this study |
| pBAD-TOPO-*dgkA* | *dgkA* under pBAD promoter in pBAD TOPO® TA | this study |
| pBAD-TOPO-*eptA-4* | *eptA-4* under pBAD promoter in pBAD TOPO® TA | this study |

**Table S6. List of primers used in this study**

| Name | Sequence 5'-3' | Description | DNA matrix |
|---|---|---|---|
| 240423-1 | GTATCGATAAGCTTGATATCGAATTCCCTTCTCTAATGCCCTTACG | *rst*A gene deletion-1 | *Vibrio harveyi* Th15_F5_F11 |
| 240423-2 | GACGATCCTAAACTGCAAGCCCCTGATGCTTGGAATGCGTAATG | *rst*A gene deletion-2 | *Vibrio harveyi* Th15_F5_F11 |
| 240423-3 | CATTACGCATTCCAAGCATCAGGGGCTTGCAGTTTAGGATCGTC | *rst*A gene deletion-3 | *Vibrio harveyi* Th15_F5_F11 |
| 240423-4 | CCCCCGGGCTGCAGGAATTCTGTTTCGTCTGTTCTGGTGC | *rst*A gene deletion-4 | *Vibrio harveyi* Th15_F5_F11 |
| 120523-1 | GTGAGCGGATAACAAAGGAAGGGCCCATGAATGTACAAGACACTAAGC | *rst*A complementation (forward) | *Vibrio harveyi* Th15_F5_F11 |
| 120523-2 | CGACGCGTCTGCAGCTCGAGTTACGCATTCCAAGCATCAG | *rst*A complementation (reverse) | *Vibrio harveyi* Th15_F5_F11 |
| 240423-10 | AGTGCACCTGCGCCAGCACC | PCR *rst*A (forward) | *Vibrio harveyi* Th15_F5_F11 |
| 240423-11 | AGGCATCGCAACTAAACAGC | PCR *rst*A (reverse) | *Vibrio harveyi* Th15_F5_F11 |
| V1-dgkA-eptA-F | CACCAGGAGATAATAATTGAAACCAGGTAAAACAGG | *dgk*A cloning (forward) | *Vibrio owensii* Th15_Z_G08 |
| V1-dgkA-R | CTAAAGTAAAATCGACGCCCAAACAAATAGGG | *dgk*A cloning (reverse) | *Vibrio owensii* Th15_Z_G08 |
| V1-eptA-F | CACCAGGAGATAATAAATGAAGACAATTGAACTCCC | *ept*A-1 cloning (forward) | *Vibrio owensii* Th15_Z_G08 |
| V1-dgkA-eptA-R | TTAGCTTTGACGACAAGTCTTAAAGATATCTAAAGCG | *ept*A-1 cloning (reverse) | *Vibrio owensii* Th15_Z_G08 |
| V4-eptA-F | CACCAGGAGATAATAAATGAATCTACCACGCTTAAAC | *ept*A-4 cloning (forward) | *Vibrio splendidus* 7T7-2 |
| V4-eptA-R | CTACCGACACGCAGCAAAAATGTC | *ept*A-4 cloning (reverse) | *Vibrio splendidus* 7T7-3 |
| dgkA_32F | GGCTCAGCAGTATAGGTCCC | *dgk*A RT-qPCR (forward) | *Vibrio harveyi* Th15_F5_F11 |
| dgkA_94R | CGTCCAATATCCGGCGAATG | *dgk*A RT-qPCR (reverse) | *Vibrio harveyi* Th15_F5_F11 |
| eptA_175F | TTTGCAGCGCTGAACTTCTT | *ept*A-1 RT-qPCR (forward) | *Vibrio harveyi* Th15_F5_F11 |
| eptA_296R | AGTGTGCCGTAGTTGTAGCT | *ept*A-1 RT-qPCR (reverse) | *Vibrio harveyi* Th15_F5_F11 |
| VH_2913_640F | AAGAAGCACGCAATCATCGC | VH_2913 RT-qPCR (forward) | *Vibrio harveyi* Th15_F5_F11 |
| VH_2913_750R | GTGACCAAGAACCGTTGCAC | VH_2913 RT-qPCR (reverse) | *Vibrio harveyi* Th15_F5_F11 |
| VH_0852_53F | CCCATGCAGCGATTGAAGTG | VH_0852 RT-qPCR (forward) | *Vibrio harveyi* Th15_F5_F11 |
| VH_0852_185R | TGAGCCAATTCTGCATTCGA | VH_0852 RT-qPCR (reverse) | *Vibrio harveyi* Th15_F5_F11 |
